# Copy-scAT: Deconvoluting single-cell chromatin accessibility of genetic subclones in cancer

**DOI:** 10.1101/2020.09.21.305516

**Authors:** Ana Nikolic, Divya Singhal, Katrina Ellestad, Michael Johnston, Aaron Gillmor, Sorana Morrissy, Jennifer A Chan, Paola Neri, Nizar Bahlis, Marco Gallo

## Abstract

Single-cell epigenomic assays have tremendous potential to illuminate mechanisms of transcriptional control in functionally diverse cancer cell populations. However, application of these techniques to clinical tumor specimens has been hampered by the current inability to distinguish malignant from non-malignant cells, which potently confounds data analysis and interpretation. Here we describe Copy-scAT, an R package that uses single-cell epigenomic data to infer copy number variants (CNVs) that define cancer cells. Copy-scAT enables studies of subclonal chromatin dynamics in complex tumors like glioblastoma. By deploying Copy-scAT, we uncovered potent influences of genetics on chromatin accessibility profiles in individual subclones. Consequently, some genetic subclones were predisposed to acquire stem-like or more differentiated molecular phenotypes, reminiscent of developmental paradigms. Copy-scAT is ideal for studies of the relationships between genetics and epigenetics in malignancies with high levels of intratumoral heterogeneity and to investigate how cancer cells interface with their microenvironment.

## INTRODUCTION

Single-cell genomic technologies have made enormous contributions to the deconvolution of complex cellular systems, including cancer (*1*). Single-cell RNA sequencing (scRNA-seq) in particular has been widely employed to understand the implications of intratumoral transcriptional heterogeneity for tumor growth, response to therapy and patient prognosis (*2*–*6*). This field has hugely benefited from an emerging ecosystem of computational tools that have enabled complex analyses of scRNA data. Since copy number variants (CNVs) mostly accrue in malignant cells and are rare in non-malignant tissues, computational platforms that use scRNA data to call CNVs have resulted in improved understanding of the behavior of genetic subclones in tumors (*7*–*9*).

On the other hand, the application of single-cell epigenomic techniques, including the assay for transposase accessible chromatin (scATAC) (*10, 11*), to study cancer has been slowed by computational bottlenecks. For instance, unlike scRNA-seq, currently no dedicated tool exists to call CNVs using scATAC data. This technical gap has prevented scATAC studies of clinical tumor specimens, which often are surgical resections that include both malignant and non-malignant cells. Inability to deconvolute these cell populations after the generation of scATAC datasets would confound downstream analyses and interpretation of this data type.

In this report, we describe Copy-scAT (Copy number inference using scATAC-seq data), a new computational tool that uses scATAC datasets to call CNVs at the single-cell level. Using scATAC datasets from adult glioblastoma (aGBM), pediatric GBM (pGBM) and multiple myeloma (MM), we demonstrate the effectiveness of Copy-scAT in calling (A) focal amplifications, (B) segmental gains and losses and (C) chromosome arm-level gains and losses. At the most basic level, Copy-scAT can therefore discriminate between malignant and non-malignant cells in scATAC datasets based on the presence or absence, respectively, of CNVs. This distinction is fundamental to ensure that downstream analyses include only the appropriate tumor or microenvironment cell populations. At a more sophisticated level, we show that implementation of Copy-scAT allows investigations of the relationship between genetic and epigenetic principles governing the behavior of individual subclones. In this regard, we show that each genetic subclone has characteristic accessible chromatin profiles, indicating that genetics imparts information that determines key epigenetic features. Strong influence of genetics on chromatin states is demonstrated by the predisposition of genetic subclones to have stem-like or more differentiated molecular profiles in GBM.

## RESULTS

### Design and implementation of Copy-scAT

We designed Copy-scAT, an R package that uses scATAC-seq information to infer copy number alterations. Copy-scAT uses fragment files generated by cellranger-atac (10xGenomics) as input to generate chromatin accessibility pileups, keeping only barcodes with a minimum number of fragments (defaulting to 5,000 fragments). It then generates a pileup of total coverage (number of reads × read lengths) over bins of determined length (1 million bp as default) **(Fig. 1a)**. Binned read counts then undergo linear normalization over the total signal in each cell to account for differences in read depth, and chromosomal bins which consist predominantly of zeros (at least 80% zero values) are discarded from further analysis. All parameters, including reference genome, bin size, and minimum length cut-off are user-customizable. Copy-scAT then implements different algorithms to detect focal amplifications and larger-scale copy number variation.

**Fig 1.**
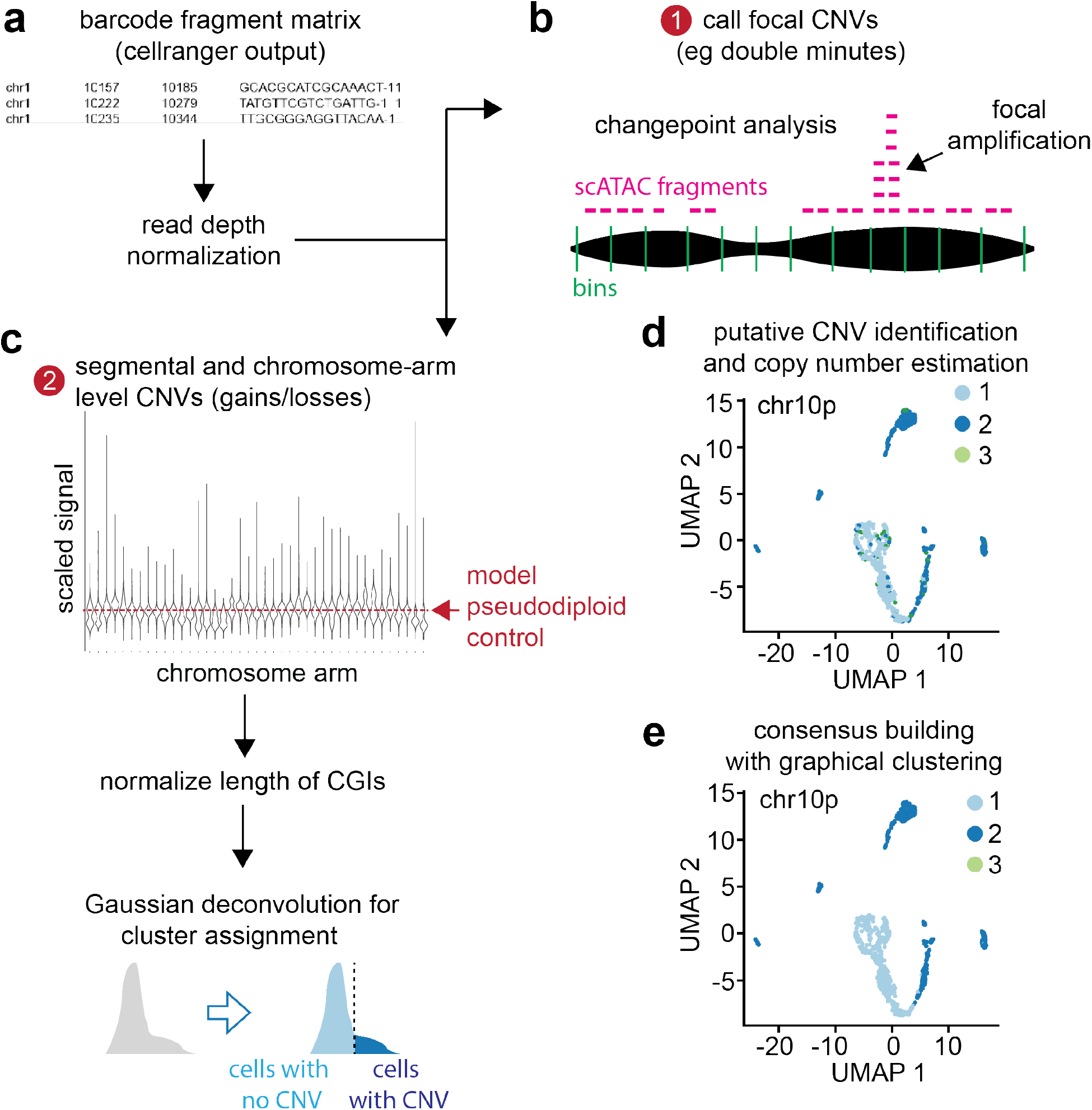
Copy-scAT workflow. **(a)** Copy-scAT accepts barcode fragment matrices generated by cellranger (10xGenomics) as input. **(b)** Large peaks in normalized coverage matrices can be used to infer focal CNVs. **(c)** Normalized matrices can be used to infer segmental and chromosome-arm level CNVs. **(d)** Example of chromosome-arm level CNV (chromosome 10p loss) called by Copy-scAT **(e)** Consensus clustering is used to finalize cell assignment.

To call focal amplifications **(Fig. 1b)**, Copy-scAT generates a linear scaled profile of density over normalized 1 Mbp bins along each chromosome on a single-cell basis, centering on the median and scaling using the range. Copy-scAT then uses changepoint analysis (*12*) (see Methods) to identify segments of abnormally high signal (Z score > 5) along each chromosome in each single cell. These calls are then pooled together to generate consensus regions of amplification, in order to identify putative double minutes and extrachromosomal amplifications. Each cell is scored as positive or negative for each amplified genomic region.

Segmental losses are called in a similar fashion, by calculating a quantile for each bin on a chromosome, running changepoint analysis to identify regions with abnormally low average signal, and then using Gaussian decomposition of total signal in that region to identify distinct clusters of cells.

For larger copy number alterations, Copy-scAT pools the bins further at the chromosome arm level using a trimmed mean, while normalizing the data on the basis of length of CpG islands contained in each bin **(Fig. 1c)**. Data is then scaled for each chromosome arm, compared to a pseudodiploid control (expected signal distribution for a diploid genotype) that is modeled for each sample, and cluster assignments are generated using Gaussian decomposition. Cluster assignments are then normalized to get an estimate of copy number for each cell **(Fig. 1d)**. These assignments can be optionally combined with clustering information to generate consensus genotypes for each cluster of cells and further filter false positives **(Fig. 1e)** For full details regarding the execution of Copy-scAT, see Methods. A step-by-step tutorial for Copy-scAT is available on GitHub (see Methods).

### Copy-scAT effectively calls CNVs in diverse malignancies

We have tested the ability of Copy-scAT to use scATAC data to call CNVs with three different approaches and with different tumor types. First, we benchmarked Copy-scAT against CNV calls made with whole-genome sequencing (WGS) data for adult GBM (aGBM) surgical resections (n = 4 samples, 3,647 cells). This approach consisted in isolating nuclei from flash-frozen aGBM samples, mixing nuclei in suspension, and then using these nuclei for either scATAC or WGS library construction **(Fig. 2a)**. This was meant to ensure similar representation of genetic subclones, which are usually regionally contiguous in this solid tumor, in both scATAC and WGS libraries. Second, we benchmarked Copy-scAT against CNV calls made using pediatric GBM (pGBM) surgical resections (n = 6 patient-matched diagnostic-relapse samples, 33,695 cells). In this case, scATAC and WGS libraries were generated from separate geographical regions of the same tumor **(Fig. 2b)**. Third, we benchmarked Copy-scAT against CNV calls made with the single-cell CNV (scCNV) assay (10xGenomics) using multiple myeloma (MM) clinical samples (n = 10 samples, 31,266 cells). Overall, we observed that Copy-scAT correctly inferred all or most of the CNVs that were called with WGS **(Figs. 2a**,**b; Figs. S1, S2)** or scCNV data **(Fig. 2c; Fig S3)**. In total, we profiled 51,571 cells from 20 malignancies from 17 patients, and were able to infer CNV status for a total of 39,486 cells **(Table S1)**. On average, we were able to call CNVs for 78.09% of cells in each sample (range: 29.16 – 91.22%) **(Table S1)**. For chromosome-arm level CNV gains, sensitivity ranged from 0.51 for MM to 1.0 for aGBM and specificity ranged from 0.93 to 0.94 **(Table S2)**. For chromosome arm-level losses, sensitivity ranged from 0.67 to 0.79 and specificity from 0.89 to 0.95. The sensitivity and specificity of focal amplifications were very high (>0.975, **Table S2**). The variation observed may reflect technical differences between the strategies used for benchmarking. As expected, the calls of Copy-scAT for aGBM were the most accurate, likely because scATAC and WGS datasets were generated by relatively homogeneous starting material, as described above. Because of its design, it is also possible that Copy-scAT is more sensitive at inferring CNVs that occur in relatively rare subclones compared to WGS, potentially explaining (in addition to true false positives) why the number of CNVs inferred by our new tool is sometime higher than inferences made with WGS.

**Fig 2.**
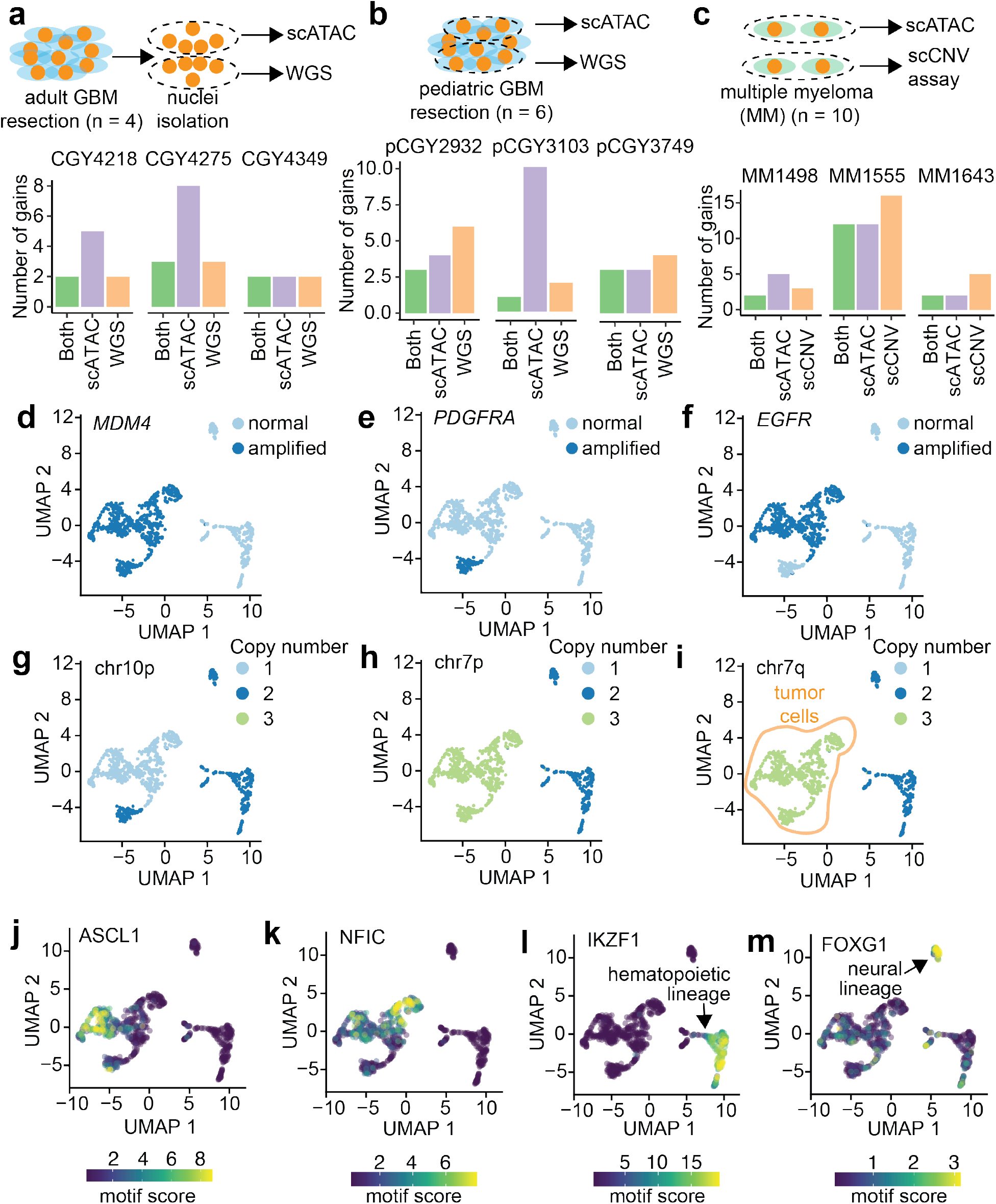
Benchmarking of Copy-scAT with three methods involving clinical samples from three distinct malignancies. **(a)** Banked frozen aGBM samples were used for both scATAC and WGS. Nuclei were isolated from the samples, mixed, and used for both scATAC and WGS. Number of chromosome-arm level gains detected in adult GBM samples identified using both methods, versus total numbers of gains detected by scATAC or WGS. **(b)** Surgical pGBM resections were split, and one section was used for scATAC and the other for WGS. Number of chromosome-arm level gains detected in adult GBM samples identified using both methods, versus total numbers of gains detected by scATAC or WGS. **(c)** Multiple myeloma samples were profiled by both scATAC and the single-cell CNV assay. Number of chromosome-arm level gains detected in adult GBM samples identified using both methods, versus total numbers of gains detected by scATAC or scCNV assay. **(d)** *MDM4* amplification in an adult GBM sample (CGY4349). Amplified cells are coloured dark blue, and normal cells in pale blue. **(e)** *PDGFRA* amplification in an adult GBM sample (CGY4349). Amplified cells are coloured dark blue, and normal cells in pale blue. **(f)** *EGFR* amplification in an adult GBM sample (CGY4349). Amplified cells are coloured dark blue, and normal cells in pale blue. **(g)** Chromosome 10p loss in an adult GBM sample. **(h)** Chromosome 7P gain in an adult GBM sample. **(i)** Chromosome 7Q gain in an adult GBM sample. **(j)** ChromVAR activity score for ASCL1. **(k)** ChromVAR activity score for NFIC. **(l)** ChromVAR activity score for IKZF1. **(m)** ChromVAR activity score for FOXG1.

### scATAC data can be used to distinguish malignant from non-malignant cells

Tumor cells often harbor CNVs, and we reasoned that the use of Copy-scAT should enable the use of scATAC data to infer CNVs and therefore to distinguish between malignant and non-malignant cells. To test this hypothesis, we overlayed CNVs called by Copy-scAT onto scATAC datasets displayed in uniform manifold approximation and projections (UMAP) plots. This exercise led to the identification of cells that were clearly positive for multiple CNVs and others that appeared to have a normal genome. As an illustrative example, we found that the aGBM sample CGY4349 was composed of discrete cell populations that harbored focal amplifications at the *MDM4* **(Fig. 2d)**, *PDGFRA* **(Fig. 2e)** and *EGFR* **(Fig. 2f)** loci, as well as chromosome 10p deletion **(Fig. 2g)** and chromosome 7 gain **(Fig. 2h**,**i)**. Copy-scAT results suggest specific lineage relationships between subclones. For instance, chromosome 7 amplifications are clonal in this sample **(Fig. 2h**,**i)**, whereas the chromosome 10 deletion is subclonal **(Fig. 2g)**. In addition, our computational tool predicts that *PDGFRA* **(Fig. 2e)** and *EGFR* **(Fig. 2f)** focal amplifications are mutually exclusive, a phenomenon that has been reported in aGBM (*13*).

Altogether, these results illustrate one specific population of cells (shaded green in **Fig. 2i**) that harbors several CNVs and are therefore putative cancer cells. At the same time, we also identified cells (labeled in dark blue in **Fig. 2i**) that did not appear to have any CNVs and are therefore likely to be cells from the tumor microenvironment. Equivalent results were obtained for pGBM **(Fig. S4)** and MM samples **(Fig. S5)**. Since the latter appear as multiple scATAC clusters, it is possible that our strategy detects multiple distinct non-neoplastic cell clusters. Differential motif analysis with ChromVAR confirmed high scores for neural progenitor cell-associated motifs like NFIC and ASCL1 in CNV^+^ cells **(Fig. 2j**,**k)**, while the putative non-neoplastic clusters showed increased occupancy at transcription factor motifs associated with hematopoietic lineages, such as IKZF1 **(Fig. 2l)**. Another CNV^-^ cluster showed enrichment of FOXG1 binding motifs in accessible chromatin, in keeping with a non-neoplastic neural cell identity **(Fig. 2)**. Using this approach, it was possible to discriminate between malignant and cells from the tumor microenvironment in all tumor samples analyzed **(Extended Figs. S6-S8)**. Copy-scAT therefore effectively uses scATAC data to infer CNVs, which can then be used to distinguish malignant from non-malignant cells and to infer lineage relationships between genetic subclones that coexist in a tumor.

### Subclonal genetics shapes chromatin accessibility profiles in aGBM

We noticed that in most tumors we analyzed, cells harboring a given CNV had a tendency to cluster together **(Fig. 2d-i)**. Individual clusters were in fact defined by the presence of specific CNVs **(Fig. 3a-c)**. This was an unexpected observation, because it is widely assumed that clustering of scATAC data reflects the global patterns of chromatin accessibility. One possible explanation for this observation could be that chromosomal regions affected by a CNV display imbalances in the fragment depth distribution of scATAC datasets, and that these patterns have a dominant effect on cluster assignment. Most scATAC-seq workflows rely on some variant of term-frequency inverse document frequency (TF-IDF) normalization rather than feature scaling, and this may amplify the effects of CNV-driven DNA content imbalances. For instance, it is possible that focal amplifications of the *PDGFRA* locus result in increased frequency of transposition events that are mapped to this site. A dominant effect of chromatin accessibility at this amplified locus could result in *PDGFRA*-amplified cells clustering together in UMAP representations of scATAC data **(Fig 3d**,**e)**. Indeed we found that compared to a random selection of peaks, the chromosomes which carried CNVs had significantly different numbers of peaks ranked as highly variant than chromosomes that did not have CNVs, leading to a markedly uneven distribution of top peaks (p < 2.2E-16; Chi-squared test; **Fig. S9a**) This was not seen in non-neoplastic cells, which had relatively even top fragment distribution patterns (p = 0.05472, Chi-squared test; **Fig. S9b**). To test this hypothesis, we used Copy-scAT to call CNVs in our tumor samples, then removed all peaks mapping to chromosomes predicted to harbor CNVs, and finally re-clustered all cells in each sample **(Fig. 3f)**. We found that although removing chromosomes with CNVs from our analyses changed the overall cluster structure of a sample **(Fig. 3g)**, *PDGFRA*-amplified cells still clustered close to each other **(Fig. 3h)**. In fact, our results indicate that clustering after CNV removal is more granular but overall very stable **(Fig. 3i)**. In this case, *PDGFRA*-amplified cells localized to a single cluster before removing chromosomes affected by CNVs. Following removal of CNV^+^ chromosomes and re-clustering, most *PDGFRA*-amplified cells still clustered together, with only a few cells merging into a cluster that included both amplified and non-amplified cells. Comparing the most variable peaks after chromosome CNV removal showed a distribution closer to normal, supporting the marked effect of the CNVs on the identification of variant peaks (*p* = 2.418E-8; **Fig. S9c**). Contrary to current views of cancer epigenomics, these data indicate that genetic subclones may have characteristic patterns of chromatin accessibility, and that a cell’s genetic background has significant influence on its likelihood of attaining specific epigenetic states.

**Fig 3.**
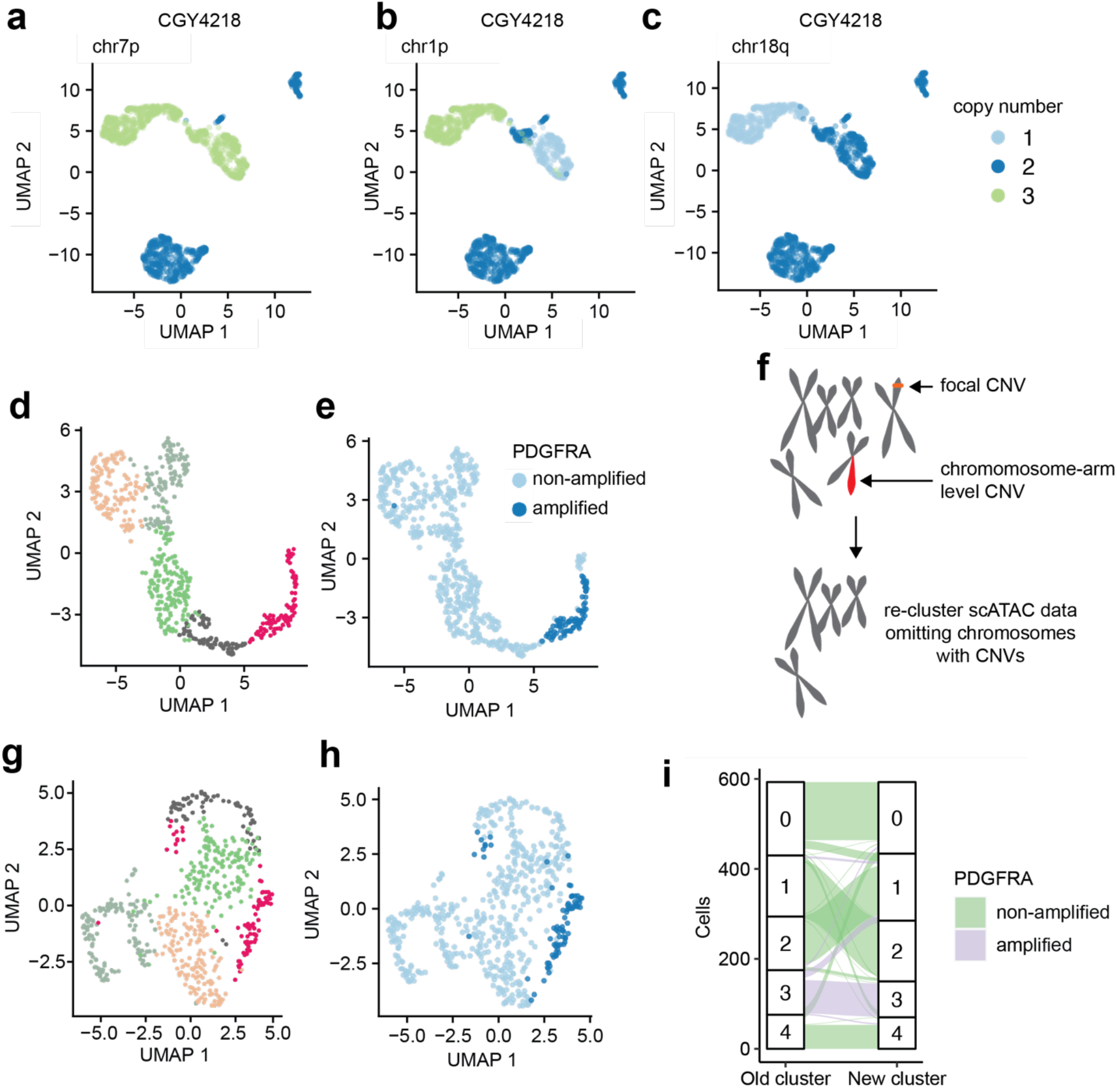
Subclonal genetics influences clustering of scATAC-seq data. **(a-c)** CNVs in adult GBM CGY4218 segregate within specific scATAC clusters. **(d, e)** *PDGFRA*-amplified cells cluster together in adult GBM CGY4349. **(f)** Diagram summarizing our strategy to remove CNVs from clustering of scATAC data. All chromosomes or regions with putative CNVs were removed from downstream analyses, and cells were re-clustered. **(g)** Reclustering of **(d)** following removal of chromosomes and regions affected by CNVs in CGY 4349. **(h)** Distribution of *PDGFRA*-amplified cells following re-clustering. **(i)** Cluster assignments of cells in CGY4349 (aGBM specimen) before and after removal of CNV-containing regions (purple: *PDGFRA*-amplified cells).

### Genetic events predispose subclones to the acquisition of developmental chromatin states

We further explored the notion that CNVs may shape chromatin accessibility profiles and its possible implications for cell fate determination. As an illustrative example, we focused on an aGBM sample (CGY4218) where CNVs at chromosome 1p characterized three genetic subclones, as determined with Copy-scAT: (i) A subclone with two copies of chromosome 1p; (ii) a subclone with loss of 1p; (iii) a subclone with gain of 1p **(Fig. 4a)**.

**Figure 4.**
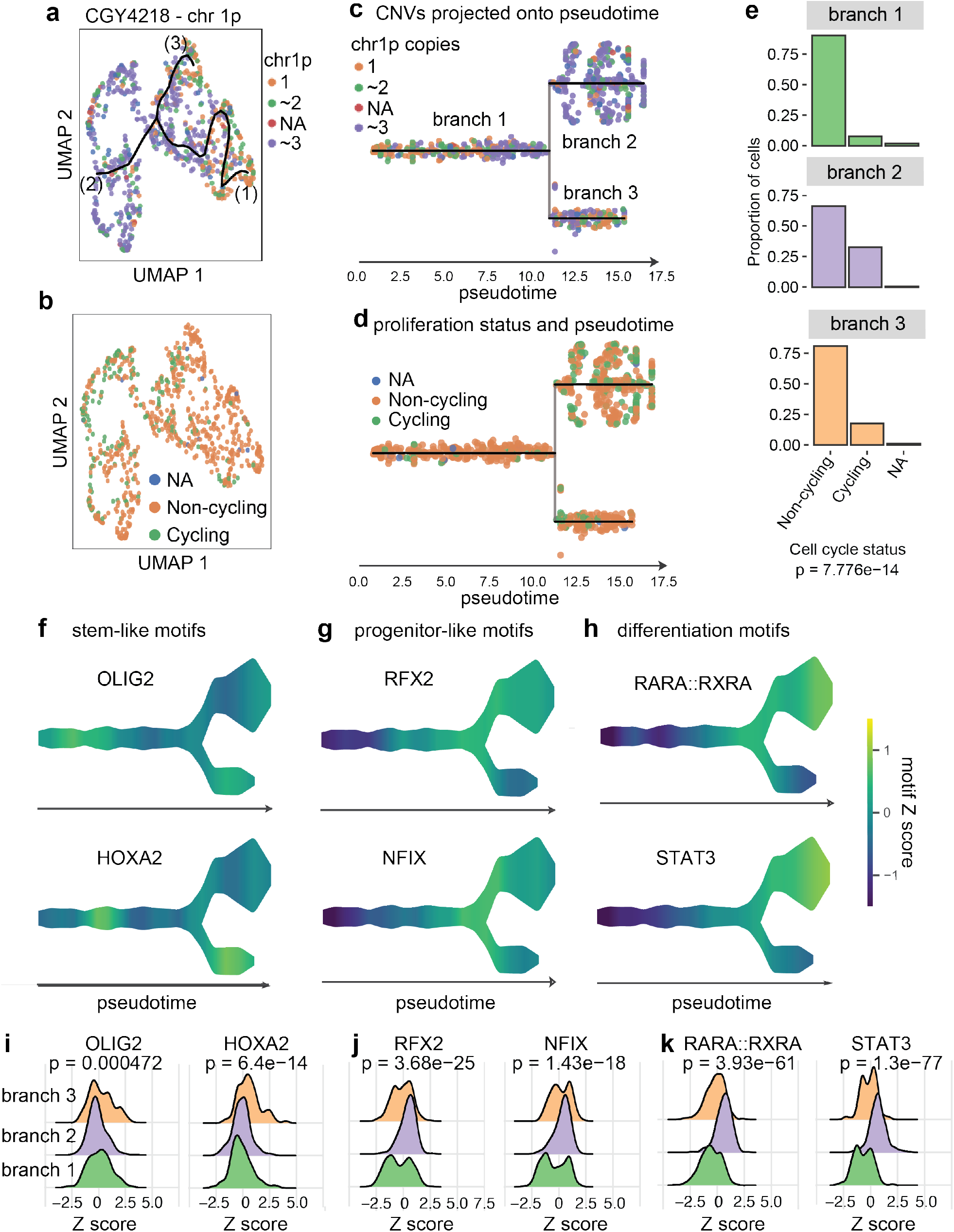
Subclonal genetic alterations predispose cells to adopt developmental chromatin states. **(a)** Cells were clustered based on scATAC ChromVAR motif scores, then shaded based on the presence of 1, 2 or 3 copies of chromosome 1P. **(b)** Cells were shaded based on their predicted cycling properties. **(a)** Data shown in **(a)** projected onto pseudotime. The resulting three branches are populated preferentially by cells with gain or loss of chromosome 1P respectively. **(d)** Proliferation status as shown in **(b)**, overlaid onto pseudotime. **(e)** Branches enriched for 1P gain show greater proportions of proliferative cells (statistics: Chi-squared test). **(f)** Scaled chromatin accessibility at binding motifs for OLIG2 and HOXA2, two TFs associated with stemness. **(g)** Scaled chromatin accessibility at binding motifs for RFX2 and NFIX, two TFs associated with progenitor-like phenotypes. **(h)** Scaled chromatin accessibility at binding motifs for RARA::RXRA and STAT3, two TFs associated with differentiated phenotypes. **(i)** Enrichment plot for motif Z scores for OLIG2 and HOXA2. **(j)** Enrichment plot for motif Z scores for RFX and NFIX. **(k)** Enrichment plot for motif Z scores for RARA::RXRA and STAT3. P values calculated by Kruskal-Wallis test.

We were interested in determining whether the major genetic subclones in this tumor had similar cycling properties. Unlike scRNA-seq, we found it is not possible to use scATAC profiles at cell cycle genes to determine whether a cell is proliferating. We reasoned that cells that are actively going through cell division have to replicate their DNA. Given that cancer cells have numerous CNVs on autosomes and could lead to noisy data, we decided to use Copy-scAT to identify cells that have doubled the number of their X chromosomes and defined them as actively cycling cells. To validate this approach, we determined the number of cells with double the number of expected X chromosomes – ie putative cycling cells – in previously published scATAC datasets for mouse brain and peripheral blood mononuclear cells (PBMCs). We hypothesized that we should be able to identify cycling cells in fetal mouse brain, but not in PBMCs. In fact, we detected numerous cycling cells (with twice the expected number of X chromosomes) in brain tissue but not in PBMCs **(Fig. S10)**. This method detected putative cycling cells in our datasets (**Fig. 4b**). We used scATAC data to arrange cells from this tumor along pseudotime with the package STREAM (*14*) **(Fig. 4c)** and then superimposed cell cycle status determined with our X chromosome doubling method **(Fig. 4d)**. The results show that cells along branch 2, which is strongly enriched for cells with chromosome 1p gains, are also the most proliferative **(Fig. 4e)**, with over 25% of the cells actively going through replication (*P* = 7.776 × 10^−14^; Chi-square test). On the other hand, ∼5% of cells along branch 1 and ∼15% of cells along branch 3 were cycling. These data therefore indicate functional differences between cells with gain or loss of chromosome 1p.

We then used ChromVAR(*15*) and STREAM-ATAC to calculate scores for transcription factor (TF) binding motifs that are associated with neurodevelopmental processes. This analysis revealed that motifs bound by TFs that are associated with stem-like phenotypes, including OLIG2 and HOXA2, are enriched in accessible chromatin regions in cells that have one copy of chromosome 1p **(Fig. 4f)**. Motifs bound by TFs associated with progenitor **(Fig 4g)** and differentiated states **(Fig. 4h)** were enriched in the branch with more cells showing gain of chromosome 1p. This was associated with a significant shift in the overall distribution of enrichment of these motifs in cells along the different branches of the trajectory (**Fig. 4i-k)**. A distribution of genetic subclones along developmental chromatin accessibility states was observed in other tumor samples we studied **(Fig. S11-S13)**. Overall, the data support the notion that tumor cells sample a discrete number of chromatin states, but their transition probabilities differ based on genotype. Consequently, chromatin states associated with each genetic subclone manifest as different functional properties, here demonstrated at the level of cell proliferation and stemness profiles.

## DISCUSSION

Here we describe Copy-scAT, the first computational tool dedicated to inferring CNVs using scATAC data. Copy-scAT resolves a computational bottleneck that has restricted the application of single-cell epigenomic techniques to the study of clinical tumor samples, which are often mixtures of malignant and non-malignant cells. The presence of non-malignant cells can severely confound the analyses of these samples and downstream data interpretation. Cell admixture is a particular problem for scATAC data because of the inherent sparsity of these datasets and because they do not provide direct information on the expression status of cell lineage markers that could be used to solve cellular identities. Because most tumor types harbor CNVs, Copy-scAT provides a simple way of solving this problem.

It is important to note that Copy-scAT enables users to perform analyses on both malignant and non-malignant cells from a tumor sample, because cell barcodes associated with both presence or absence of CNVs can be selected for downstream analyses. Implementation of Copy-scAT will therefore be beneficial to groups interested in defining the epigenomes of both tumor cells and their microenvironment. Because chromatin accessibility datasets provide information on mechanisms of transcriptional regulation by distal and proximal enhancer and super enhancer elements, Copy-scAT could be useful in clarifying epigenetic mechanisms involved in immune suppression and T cell exhaustion, for instance. Copy-scAT also allows scATAC studies of frozen banked cancer specimens (see Methods), because it requires no prior knowledge of cell composition.

We show that the underlying CNV architecture plays a significant role in clustering of scATAC data, a problem that is amplified by the use of TF-IDF algorithms for normalization. These effects are less pronounced when clustering is based on motif activity scores (e.g. ChromVAR), likely as this incorporates data from multiple chromosomes, thus dampening the effect of variation at any one specific locus. Further studies are needed to identify the optimal way to address the effects of CNVs in downstream analyses, as they may present a significant confounder and potentially mask significant biological relationships.

In this report, we provide evidence that Copy-scAT can be used to shed new light on how genetics and epigenetics interface in cancer. We show that genetic subclones tend to have unique chromatin accessibility landscapes that can promote or antagonize stem-like phenotypes. Consequently, we report that some genetic subclones have greater proportions of stem-like cells, and others appear more differentiated. These results offer a radically different view of functional hierarchies in GBM, where stem-like properties were thought to be programmed by epigenetic factors, independently of genotype. These findings provide a simple explanation for the observed intra-tumoral transcriptional heterogeneity in GBM ((*5, 16*)), by suggesting that each genetic subclone achieves specific chromatin accessibility profiles, which in turn result in subclone-specific transcriptional outcomes.

Copy-scAT will enable future studies of subclonal chromatin dynamics in complex tumor types and may be an important tool in better understanding the functional relationships between subclones, their microenvironment and therapy response.

## MATERIALS AND METHODS

### Ethics and consent statement

All samples were collected and used for research with appropriate informed consent and with approval by the Health Research Ethics Board of Alberta.

### scATAC-seq sample processing

GBM samples were either frozen surgical resections (pediatric GBM) or cells dissociated from fresh surgical specimens and cryopreserved (adult GBM). Samples were dissociated in a 1.5 mL microcentrfuge tube, using a wide-bore P1000 pipette followed by a narrow bore P1000 pipette in nuclear resuspension buffer (10 mM Tris-HCl; 10 mM NaCl; 3 mM MgCl2; 0.1% IGEPAL, 0.1% Tween-20, 0.01% Digitonin, 1% BSA in PBS), then vortexed briefly, chilled on ice for 10 minutes, then pipetted again, and spun at 4°C, 500 g for 5 minutes. This step was repeated, and the sample was then resuspended in Tween wash buffer (10 mM Tris-HCl; 10 mM NaCl; 3 mM MgCl_2_; 0.1% IGEPAL, 0.1% Tween-20; 1% BSA in PBS), then strained though a 35 μm cell strainer FACS tube (Fisher Scientific 08-771-23) to remove debris. Nuclei were then quantified by trypan blue on the Countess II (Invitrogen), spun down at 500 g at 4°C for 5 minutes, resuspended in the nuclear isolation buffer (10X Genomics), and the rest of the scATAC was performed as per the 10X Genomics protocol. MM samples were from bone marrow aspirates collected from patients; tumor cells were isolated from mononuclear cell fractions through Ficoll gradients coupled with magnetic bead sorting of CD138^+^ cells. scATAC libraries were prepared from GBM and MM samples using a Chromium controller (10xGenomics). Libraries were sequenced on NextSeq 500 or Novaseq 6000 instruments (Illumina) at the Centre for Health Genomics and Informatics (CHGI; University of Calgary) using the recommended settings.

### scATAC-seq initial data analysis

The raw sequencing data was demultiplexed using cellranger-atac mkfastq (Cell Ranger ATAC, version 1.1.0, 10x Genomics). Single cell ATAC-seq reads were aligned to the hg38 reference genome (GRCh38, version 1.1.0, 10x Genomics) and quantified using cellranger-atac count function with default parameters (Cell Ranger ATAC, version 1.1.0, 10x Genomics).

### Single-cell CNV analysis

#### Fragment pileup and normalization

The fragment file was processed and signal was binned into bins of a preset size (default 1 Mb) across the hg38 chromosomes to generate a genome-wide read-depth map. Only barcodes with a minimum of 5000 reads were retained, in order to remove spurious barcodes. This flattened barcode-fragment matrix pileup was cleaned by removal of genomic intervals which were uninformative (greater than 80% zeros) and barcodes with greater than a certain number of zero intervals. Cells passing this first filter were normalized with counts-per-million normalization using *cpm* in the *edgeR* package (*17*).

#### Chromosome arm CNV analysis

The normalized barcode-fragment matrix was collapsed to the chromosome arm level, using chromosome arm information from the UCSC (UCSC table: cytoBand), centromeres were removed, and signal in each bin was normalized using the number of basepairs in CpG islands in the interval using the UCSC CpG islands table (UCSC table: cpgIslandExtUnmasked). The signal was then summarized using a quantile-trimmed-mean (between the 50^th^ and 80^th^ quantiles). Only chromosome arms with a minimum trimmed mean signal were kept for analysis.

The chromosome arm signal matrix is mixed with a generated set proportion of pseudodiploid control cells, defined using the mean of chromosome segment medians with a defined standard deviation. This cell-signal matrix is then scaled across each chromosome arm and centered on the median signal of all chromosomes. Each chromosome arm segment is then analyzed using Gaussian decomposition with Mclust (*18*). The subsequent clusters are filtered based on Z scores and mixing proportions, and redundant clusters are combined. These Z scores are then translated into estimated copy numbers for each segment for each barcode. The barcode CNV assignments can be optionally used to assign consensus CNVs to clusters generated in other software packages such as Loupe or Seurat/Signac.

#### Detection of amplifications

The normalized barcode-fragment matrix was scaled and mean-variance changepoint analysis using the Changepoint package was performed for each cell and each chromosome to identify areas of abnormally high signal (Z score greater than 5) (*19*). The consensus coordinates of each amplification region were generated across all cells and only abnormalities affecting a minimum number of cells were kept for analysis.

#### Detection of loss of heterozygosity

The normalized barcode-fragment matrix was scaled as above. As overall coverage levels in these samples are quite sparse, a chromosome-wide coverage profile was generated for the entire sample in bulk, using the 30% quantile as a cut-off, and then changepoint analysis was used to find inflection points. This was followed by Gaussian decomposition of the values using Mclust to identify putative areas of loss or gain, thresholded by a minimum difference in signal between the clusters identified by Mclust.

### scATAC trajectory analysis

STREAM-ATAC and STREAM (*20*) were used to generate pseudotime trajectories based on motif occupancy profiles generated using ChromVAR (*21*) with the JASPAR 2018 motif database as reference (*22*). Dimensionality reduction was performed using the top 20 components and 50 neighbours, and an initial elastic graph was generated on the 2D UMAP projection using 10 clusters, using the kmeans method with n_neighbours = 30. An elastic principal graph was constructed using the parameters epg_alpha = 0.02, epg_mu = 0.05, epg_lambda = 0.02 and epg_trimmingradius = 1.2, with branch extension using ‘QuantDists’. Trees were rooted using the branch with highest motif activities for OLIG2 and ETV motifs as root.

### Whole genome sequencing

DNA was extracted from residual nuclei from the same samples and tissue fragments used for scATAC-seq of adult GBM samples, using the Qiagen DNEasy Blood and Tissue DNA extraction kit (Qiagen # 69504). Libraries were prepared using the NEBNext Ultra II DNA Library Prep Kit (#E7645) and sequenced on the Novaseq 6000 (Illumina) at the CHGI (University of Calgary), in paired-end mode.

### Whole genome data processing

Genome data was aligned to the hg38 assembly using bwa mem (bwa 0.7.17)(*23*). Samtools was used to extract high-quality reads (Q > 30) and picard tools (Broad Institute) was used to remove duplicates (*24*).

### Whole genome SNV and CNV detection

Gatk mutect2 (Broad Institute) was run on the filtered data to detect SNVs with low stringency using the following settings: *--disable-read-filter MateOnSameContigOrNoMappedMateReadFilter*. CNVkit was subsequently used to call copy number variants using the following parameters: *--filter cn -m clonal –purity 0*.*7* (*25*). Adjacent segments were further combined and averaged using bedtools (*26*).

### Data visualization and clustering

Data was visualized and UMAP plots were generated using Seurat 3.0.0 and Signac 1.0.0 (*27*) and Cell Loupe version 4.0.0 (*28*).

### Statistical analysis

Between-group differences in discrete values (e.g. chromosome peaks, branch assignments) were calculated using the Chi-squared test. Differences in non-parametric distributions (motif accessibility in clusters) were quantified using the Kruskal-Wallis test.

## Funding

A Canada Research Chair in Brain Cancer Epigenomics (tier 2) from the Government of Canada, Project grants from the Canadian Institutes of Health Research (CIHR; PJT-156278, PJT-173475), a Discovery grant from the Natural Sciences and Engineering Research Council (NSERC) and an Azrieli Future Leader in Canadian Brain Research grant to MG; a Clinician Investigator Program fellowship from Alberta Health Services and a fellowship from Alberta Innovates to AN; an Eyes High Scholarship from the University of Calgary to DS; a Clark Smith postdoctoral fellowship and a CIHR postdoctoral fellowship to MJ; a Canada Research Char in Precision Oncology (tier 2) and a CIHR grant (PJT-438802) to SM; an Alberta Graduate Excellence Scholarship and Alberta Innovates scholarship to AG. This project has been made possible by the Brain Canada Foundation through the Canada Brain Research Fund, with the financial support of Health Canada and the Azrieli Foundation.

## Author contributions

Conception and experimental design: AN, MG. Generation of datasets: AN, KE, JC, PN, NB. Data acquisition and analysis: AN, DS, MJ, AG, SM, NB, MG. Data interpretation and creation of new software: AN, MG. Manuscript preparation: All co-authors.

## Competing interests

The authors declare no competing interests.

## Data and materials availability

The Copy-scAT package and a sample tutorial are available on Github at http://github.com/spcdot/CopyscAT. All datasets will be made available upon publication in a peer reviewed journal.

## SUPPLEMENTAL MATERIAL

**Table S1.** Summary of samples and cells profiled by Copy-scAT

| Sample | Unique barcodes<br>after pileup | Unique barcodes<br>after filtering | Percent passing<br>filters |
| --- | --- | --- | --- |
| CGY4218 | 1542 | 1335 | 86.58% |
| CGY4250 | 1371 | 947 | 69.07% |
| CGY4275 | 1004 | 609 | 60.66% |
| CGY4349 | 961 | 756 | 78.67% |
| pCGY2932 | 1203 | 802 | 66.67% |
| pCGY2937 | 1445 | 1200 | 83.04% |
| pCGY3103 | 2189 | 956 | 43.67% |
| pCGY3402 | 3162 | 2318 | 73.31% |
| pCGY3749 | 2774 | 2503 | 90.23% |
| pCGY4021 | 1963 | 1382 | 70.40% |
| MM1217 | 890 | 792 | 88.99% |
| MM1388 | 2538 | 2219 | 87.43% |
| MM1389 | 7438 | 6607 | 88.83% |
| MM1438 | 1774 | 1578 | 88.95% |
| MM1460 | 2048 | 1683 | 82.18% |
| MM1479 | 5135 | 4564 | 88.88% |
| MM1498 | 7408 | 2160 | 29.16% |
| MM1555 | 7220 | 6586 | 91.22% |
| MM1643 | 3141 | 2794 | 88.95% |
| MM1698 | 2844 | 2283 | 80.27% |
| **Total cells profiled** | **58050** | **44074** | **76.86%** |

**Table S2.** Sensitivity and Specificity of Copy-scAT in aGBM, pGBM and MM samples

|  | Gains |  | Losses |  | Amplifications |  |
| --- | --- | --- | --- | --- | --- | --- |
| Samples | Sensitivity | Specificity | Sensitivity | Specificity | Sensitivity | Specificity |
| aGBM (n = 3) | 1.0 | 0.94 | 0.79 | 0.89 | 1.0 | 1.0 |
| pGBM (n= 6) | 0.73 | 0.93 | 0.73 | 0.95 | N/A | 0.975 |
| MM (n = 10) | 0.51 | 0.94 | 0.67 | 0.89 | N/A | N/A |

**Fig S1.**
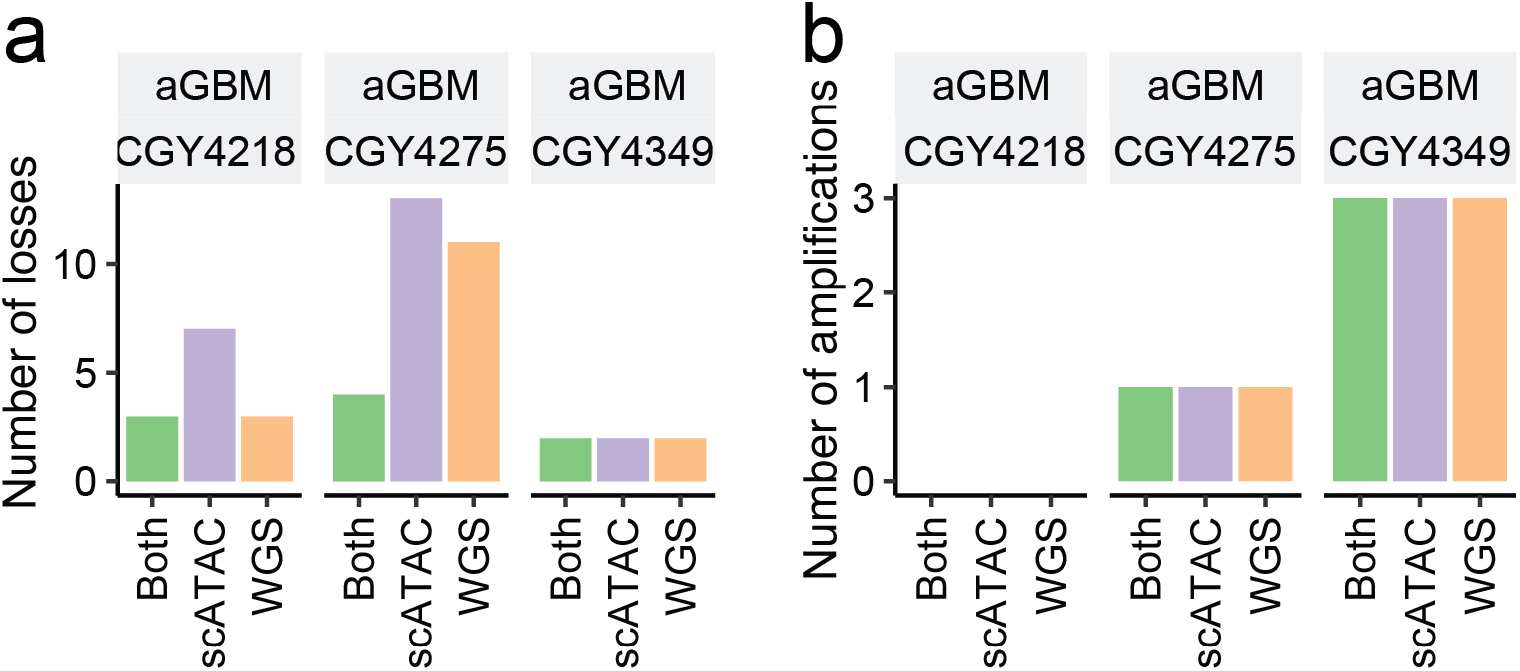
Comparison of CNVs inferred by Copy-scAT and by WGS for adult GBM samples. **(A)** Comparison of chromosome arm level losses detected in three adult GBM samples by single cell ATAC, WGS, or both methods. **(B)** Comparison of focal amplifications detected in three adult GBM sample by scATAC, WGS, or both methods.

**Fig S2.**
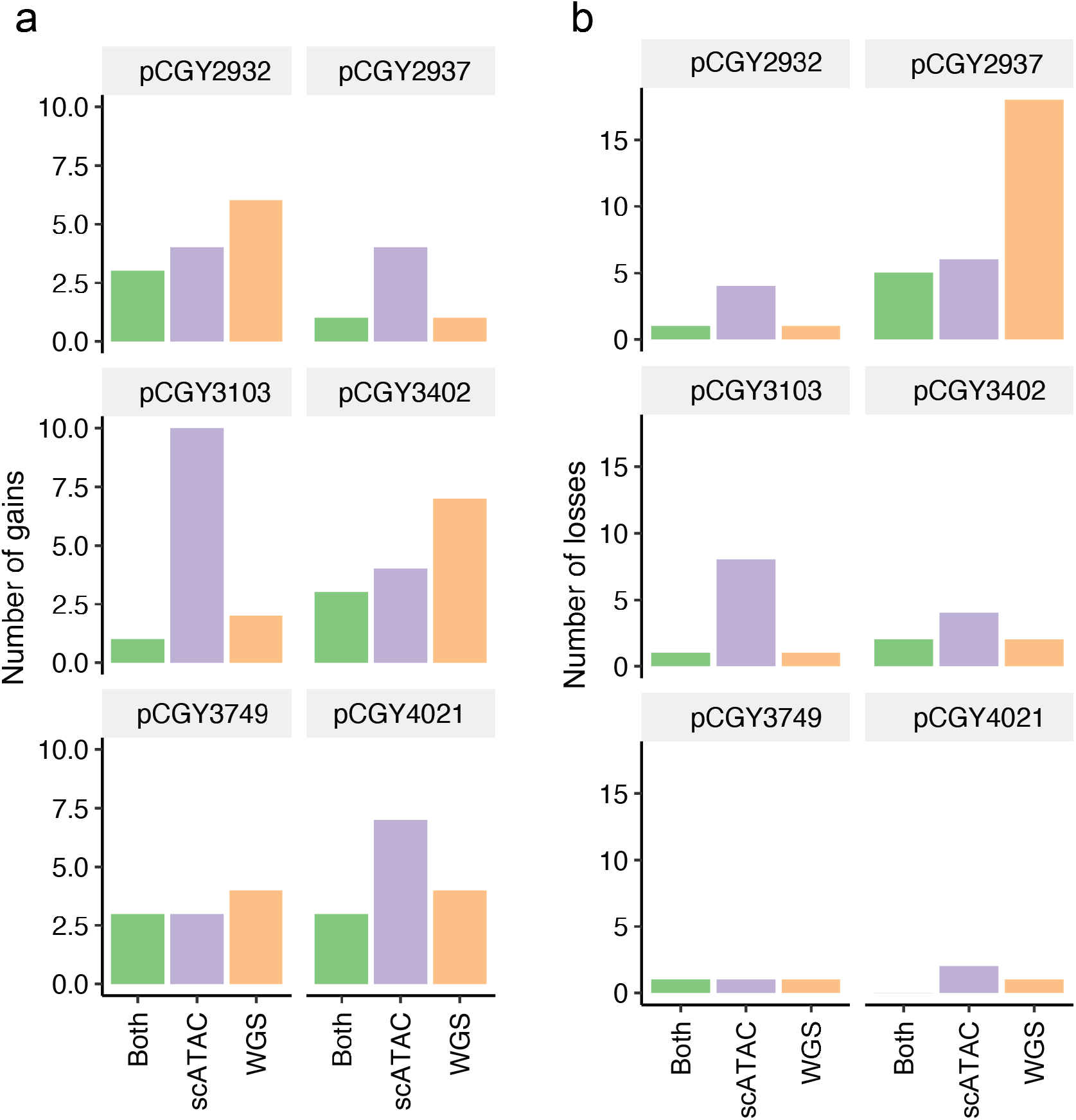
Comparison of CNVs inferred by Copy-scAT or WGS in pediatric GBM samples. **(a)** Gains detected in three pediatric GBM samples compared to linked-reads WGS. **(b)** Losses detected in three pediatric GBM samples compared to linked-reads WGS.

**Fig S3.**
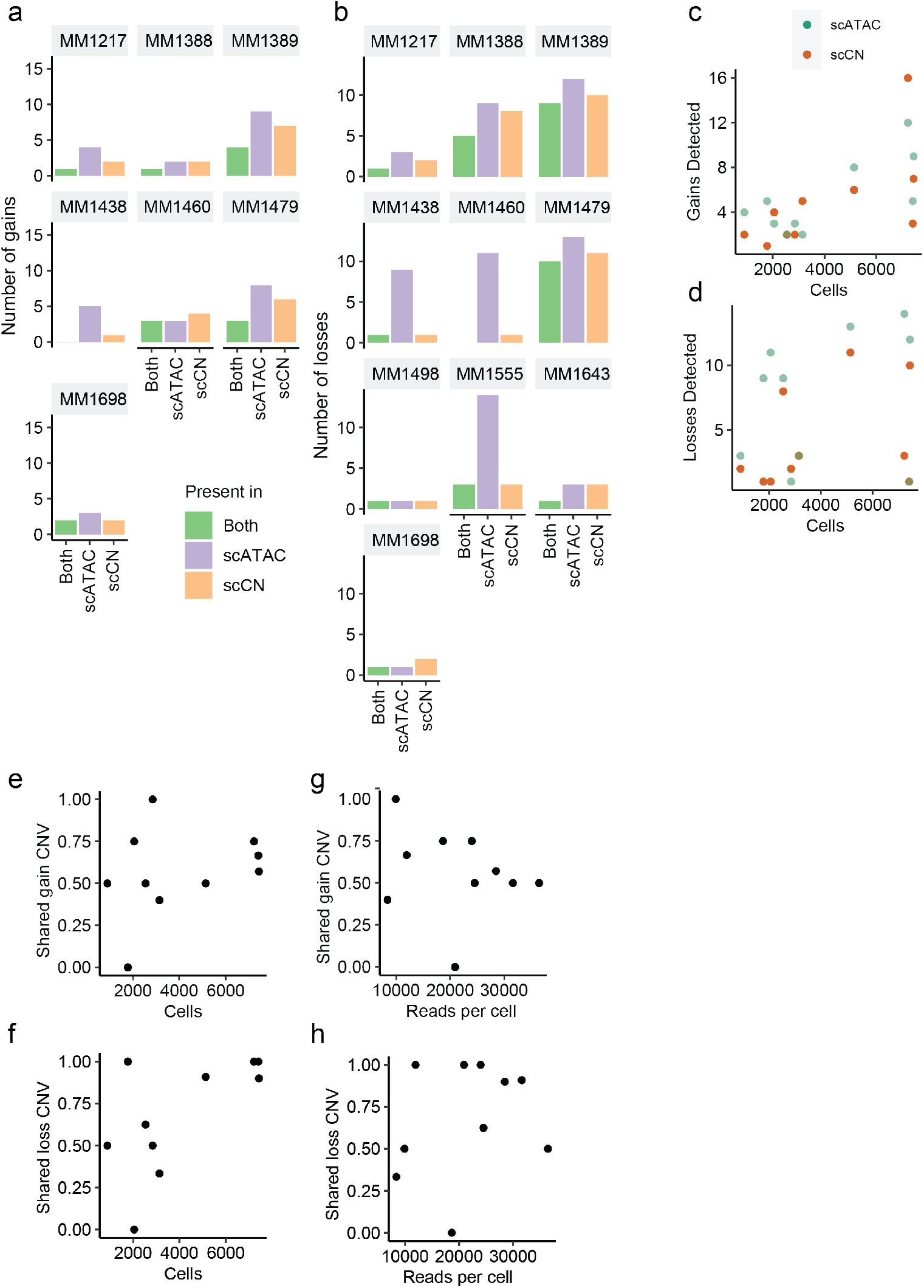
Comparison of CNVs inferred by Copy-scAT or with the scCNV assay in multiple myeloma samples. **(a)** Comparison of gains seen in additional myeloma samples versus 10x single-cell CNV sequencing. **(b)** Comparison of chromosome losses seen in additional myeloma samples versus 10x single-cell CNV sequencing. **(c**,**d)** Number of gains and losses detected by both methods compared to number of cells in scATAC-seq sample. **(e-f)** Number of shared gains or losses detected between the two methods, plotted versus the number of cells in the scATAC-seq experiment. **(g-h)** Number of shared gains or losses detected between the two methods, plotted versus the number of reads per cell in the scATAC.

**Fig S4.**
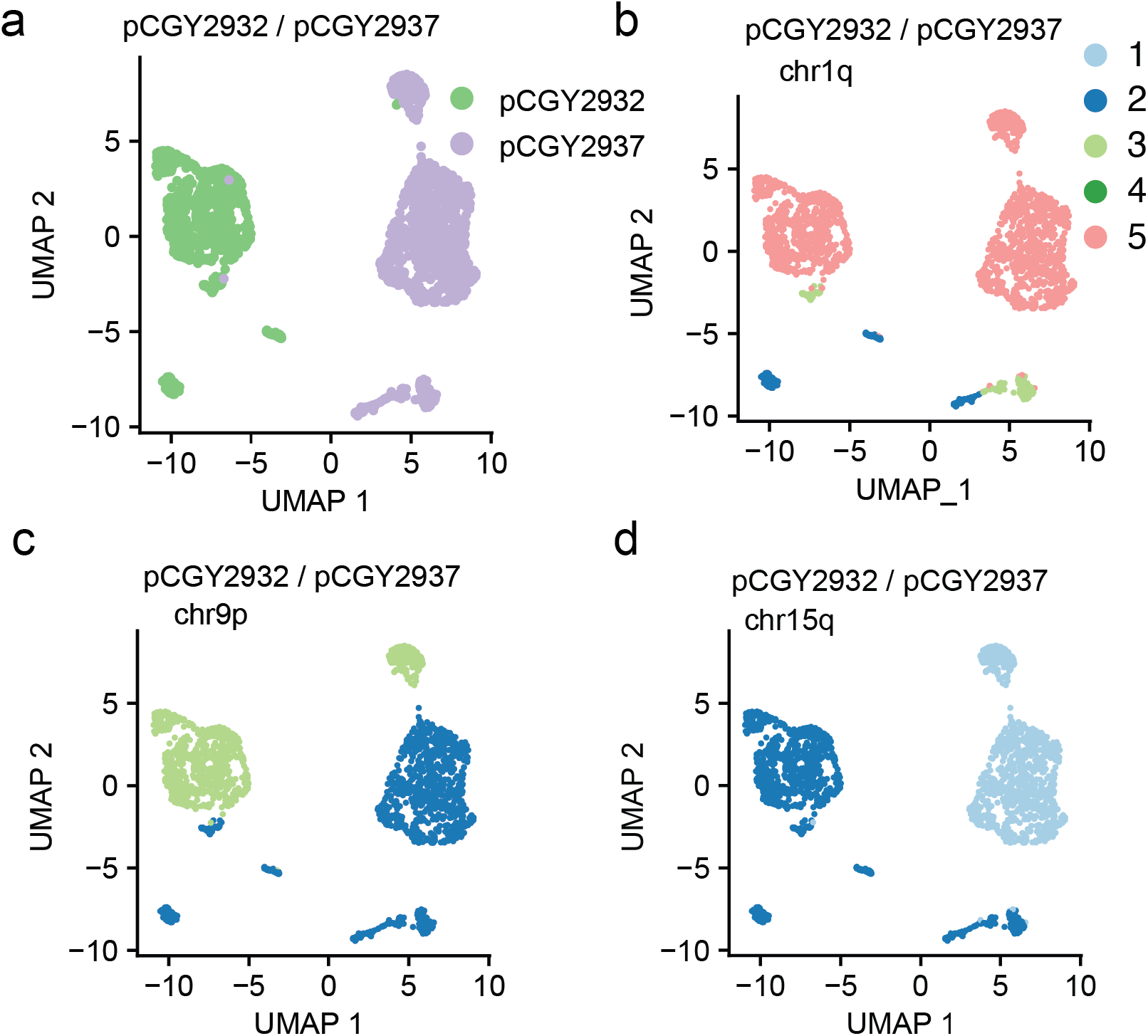
CNVs are detected in scATAC clusters with Copy-scAT in pediatric GBM samples. **(a)** Overview of cell assignments in two paired patient libraries. **(b-d)** Representative WGS-confirmed alterations detected in pCGY2932 and pCGY2937.

**Fig S5.**
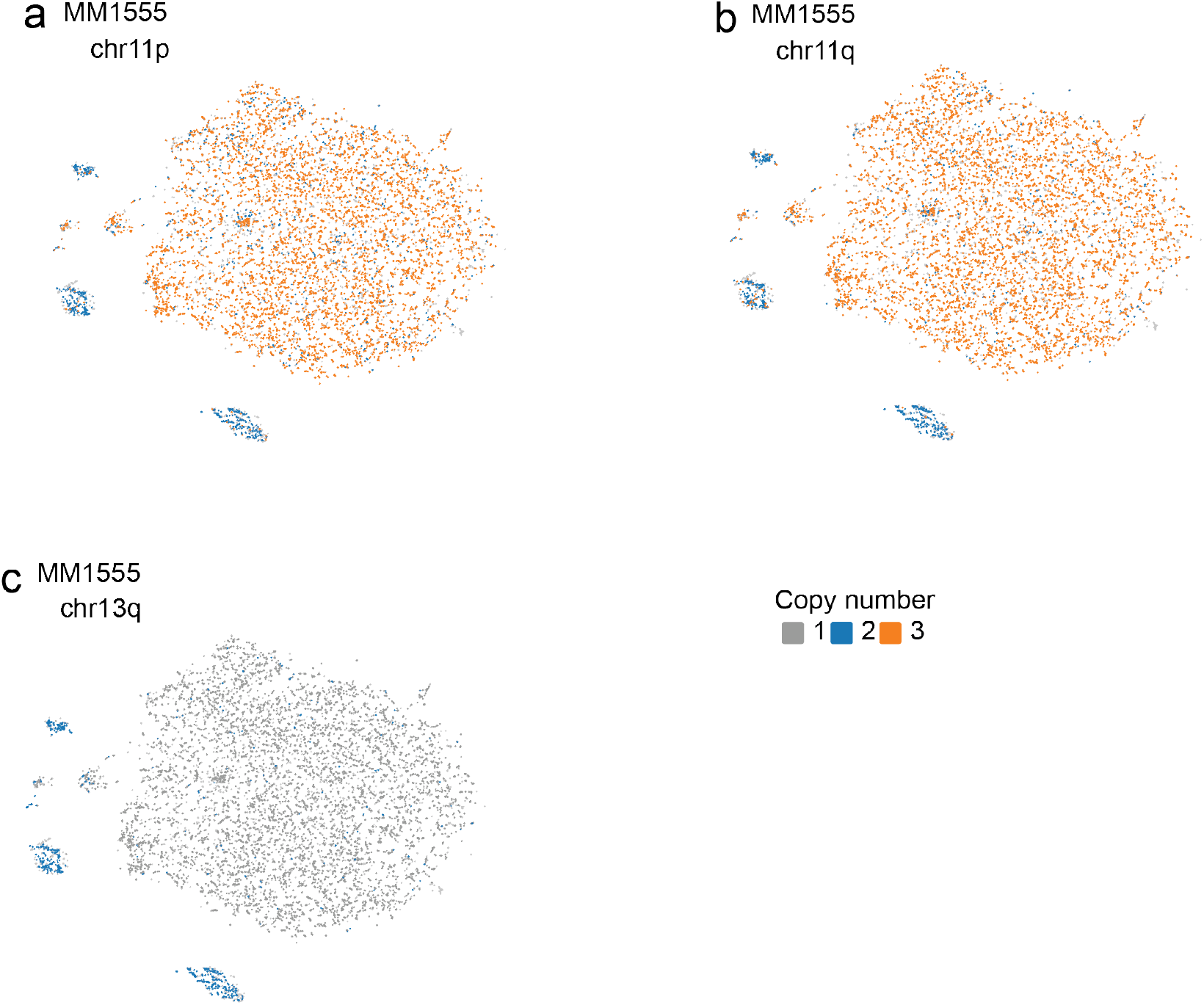
CNVs are identified by Copy-scAT in specific scATAC clusters in multiple myeloma samples. **(a)** Gain of chromosome 11p restricted to neoplastic cell populations. **(b)** Similar pattern with gain of chromosome 11q. **(c)** Similar pattern with loss of chr13q.

**Fig S6.**
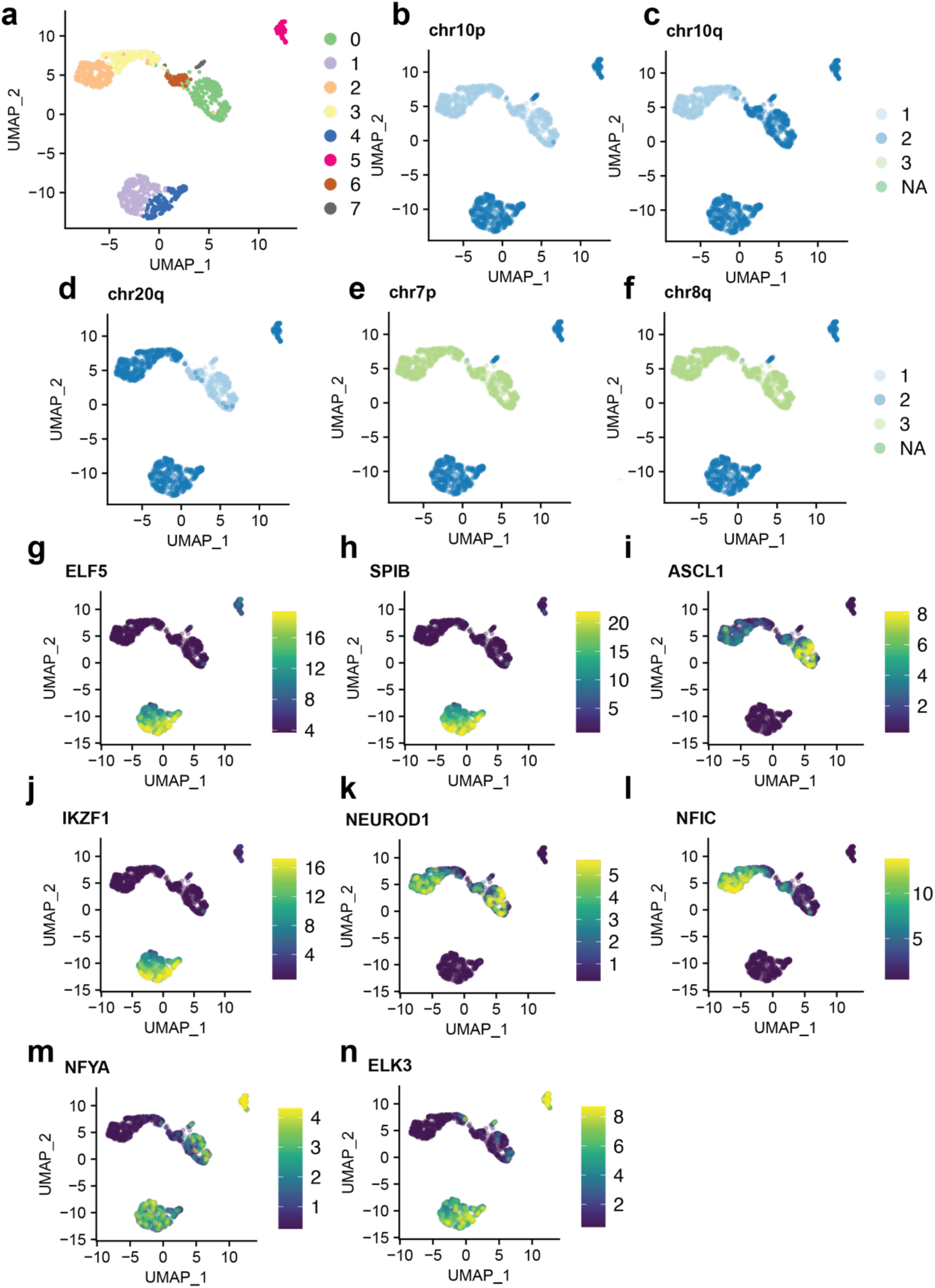
Additional chromosome copy number analyses for CGY4218. **(a)** Initial neighbourhood clustering results from Signac. **(b-f)** Representative chromosome-level copy number alteration profiles for tumour and normal cells. **(g-n)** Representative motif scores from ChromVAR for different motifs, including **(g)** ELF5, **(h)** SPIB, **(i)** ASCL1, **(j)** IKZF1, **(k)** NEUROD1, **(l)** NFIC, **(m)** NFYA, **(n)** ELK3.

**Fig S7.**
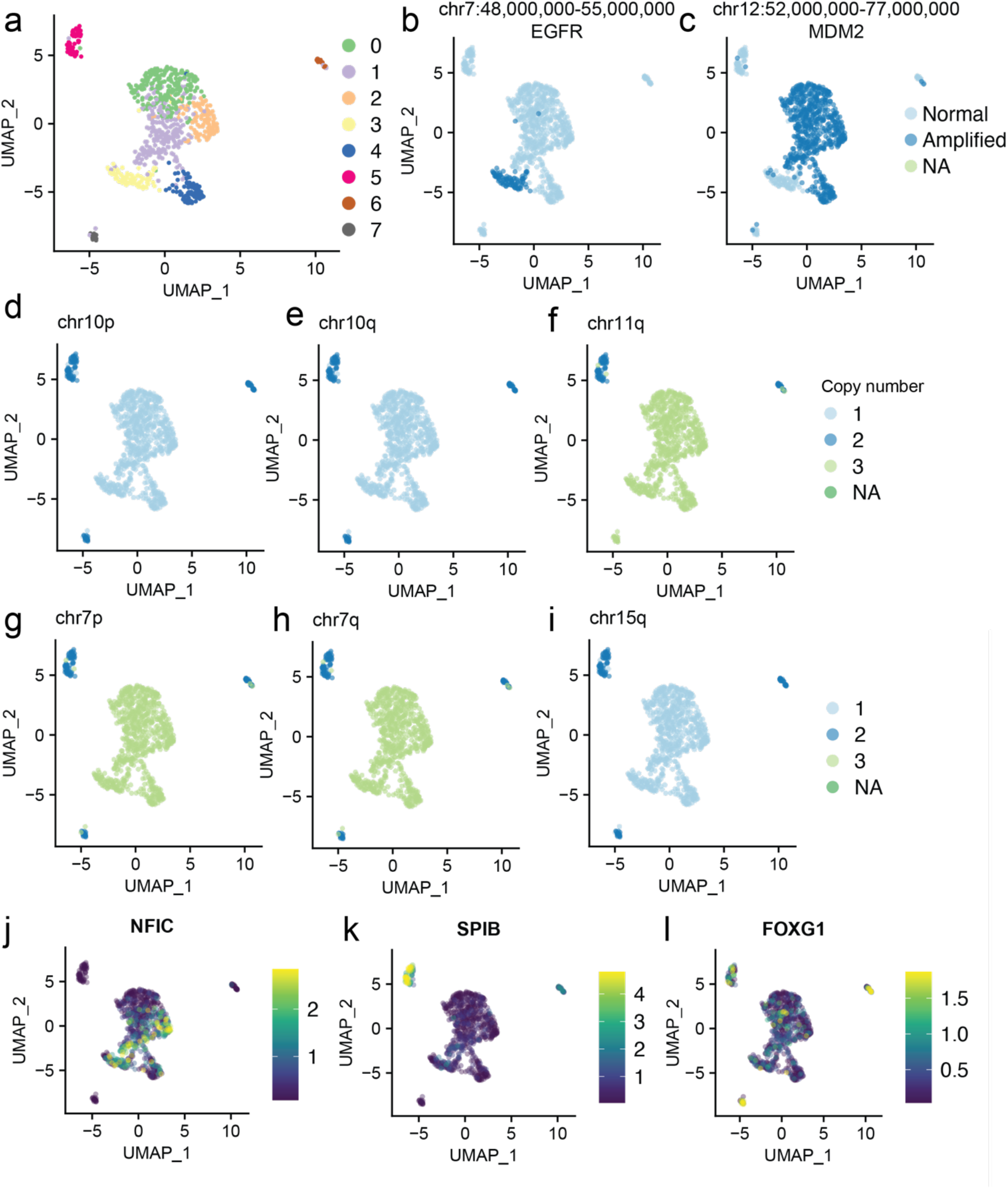
Representative copy number information and distribution for aGBM sample CGY4250. **(a)** Neighbourhood clustering results from Signac. **(b-c)** Distribution of amplifications in *EGFR* and *MDM2*. **(d-i)** Representative chromosome-level copy number alteration profiles for tumour and normal cells. **(j-l)** Representative motif scores from ChromVAR for different motifs, including **(j)** NFIC, **(k)** SPIB and **(l)** FOXG1.

**Fig S8.**
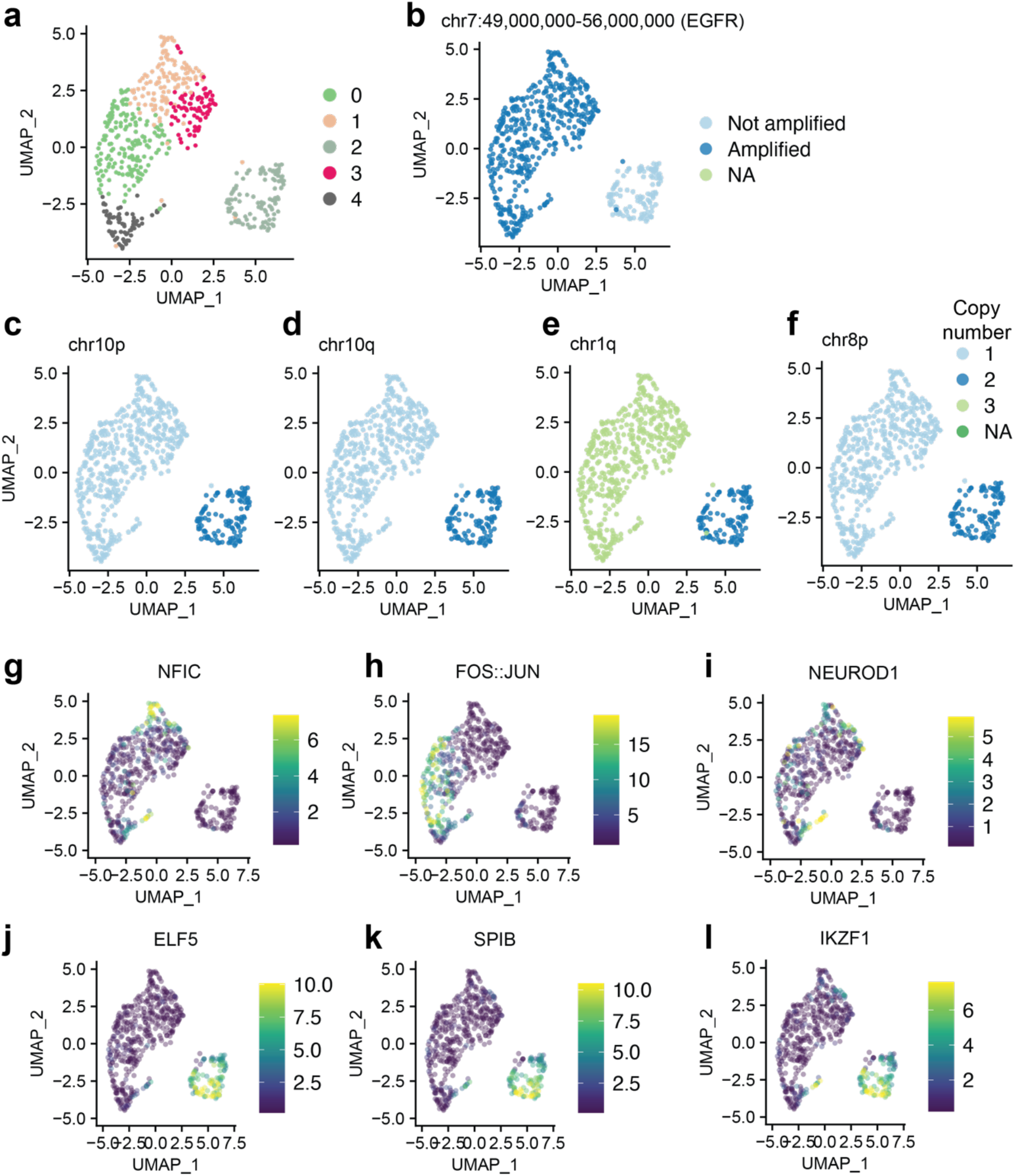
Representative copy number information and distribution for aGBM sample CGY4275. **(a)** Neighbourhood clustering results from Signac. **(b)** Distribution of amplifications in *EGFR*. **(c-j)** Representative chromosome-level copy number alteration profiles for tumour and normal cells. **(g-l)** Representative motif scores from ChromVAR for different motifs, including **(g)** NFIC, **(h)** FOS::JUN, **(i)** NEUROD1, **(j)** ELF5, **(k)** SPIB, and **(l)** IKZF1.

**Fig S9.**
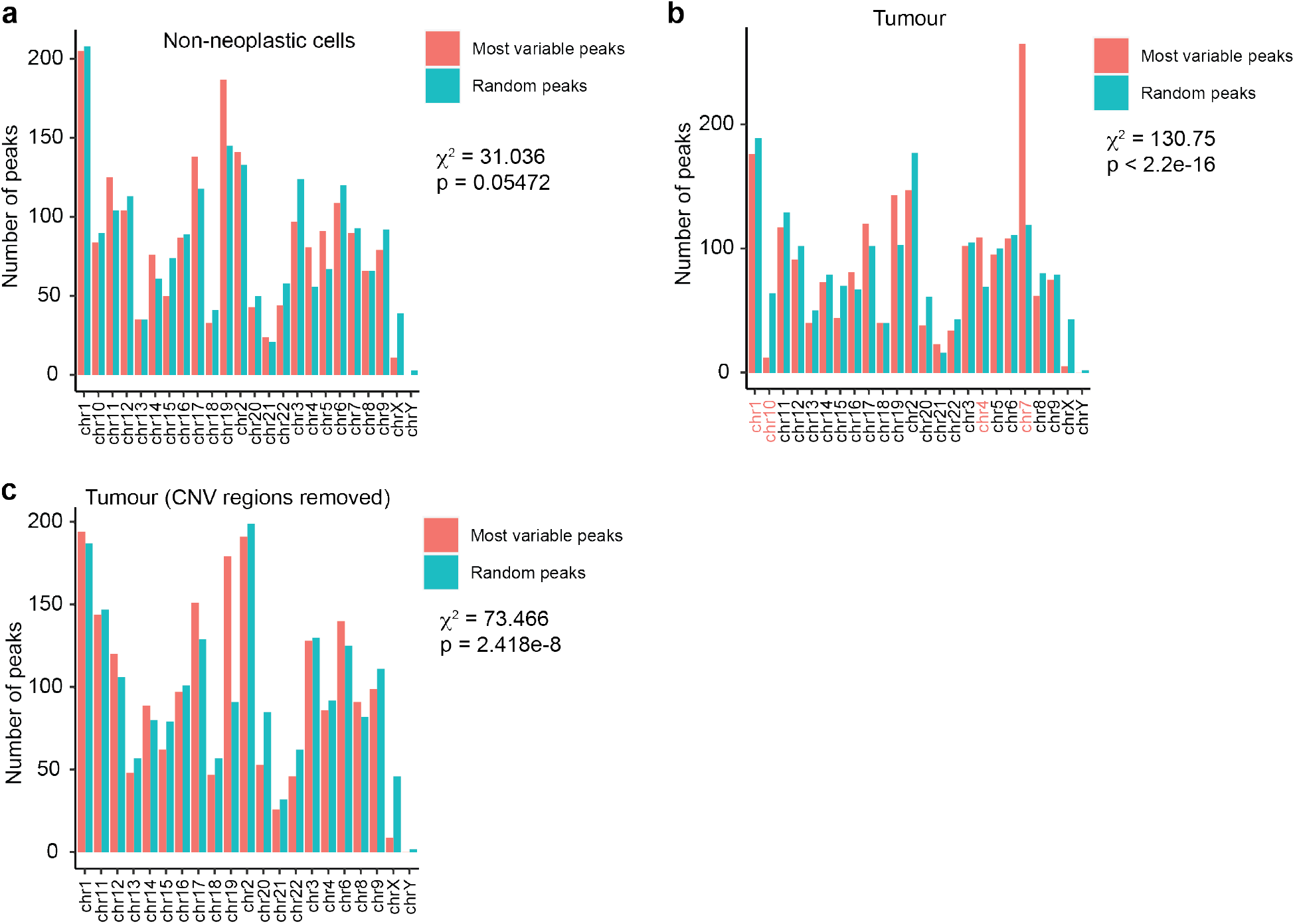
Effects of removing CNVs on variance in aGBM sample CGY4349. **(a)** Distribution of the top 2000 most variable peaks in the tumour cells after filtering out non-neoplastic cells; p value from Chi-squared test. **(b)** Distribution of top 2000 most variable peaks in non-neoplastic cells after filtering (P VALUE FROM CHI-SQUARED TEST). Chromosomes with CNVs or amplification regions are highlighted in pink. **(c)** Distribution of top 2000 most variable peaks in tumour cells after filtering of non-neoplastic cells and removal of regions containing CNVs (P VALUE FROM CHI-SQUARED TEST).

**Fig S10.**
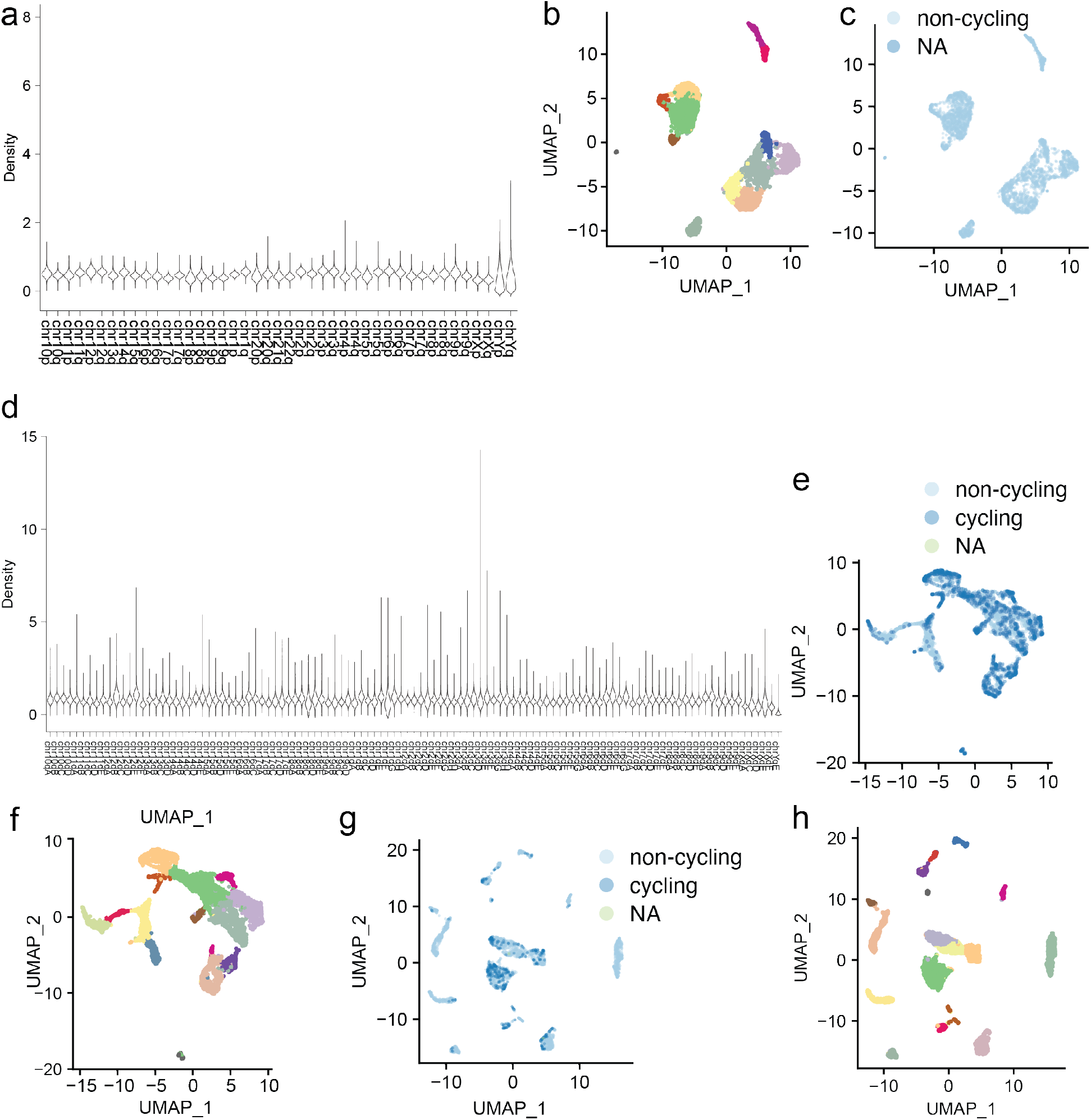
Validation of Copy-scAT and identification of putative proliferative cells in non-neoplastic datasets. **(a)** Chromosome copy number distribution in a 10X dataset of 5000 human PBMCs. **(b)** Seurat clusters for the 10X dataset of 5000 human PBMCs. **(c)** Estimate of cycle status for the 10X dataset of 5000 human PBMCs. **(d)** Chromosome copy number distribution in a 10X dataset of mouse embryonic brain at E18. **(e**,**f)** Predicted cycle status and cluster assignments in E18 mouse brain. **(g**,**h)** Predicted cell cycle status and cluster profile in P50 mouse brain dataset from 10X.

**Fig S11.**
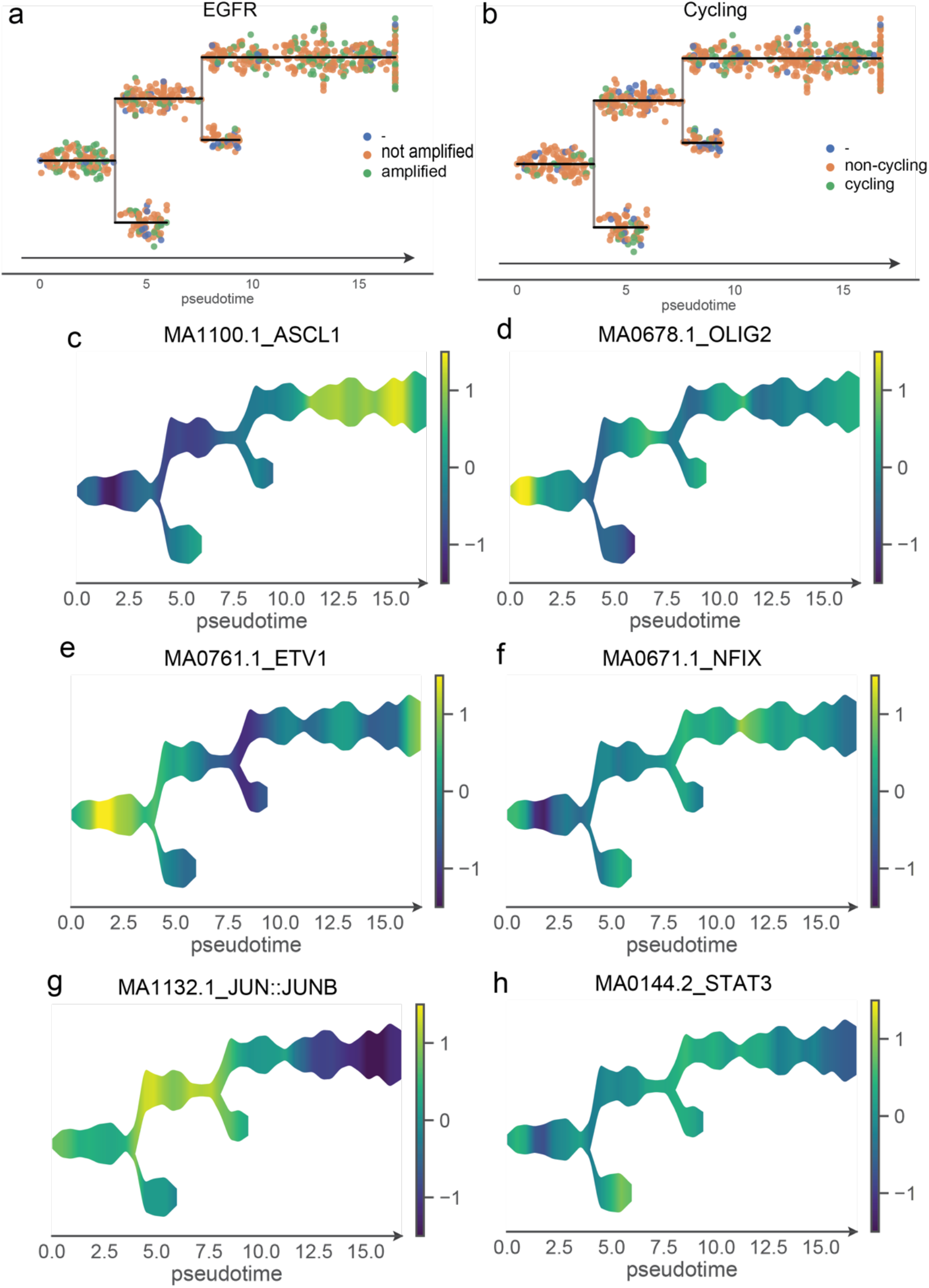
Pseudotime trajectory analysis of aGBM sample CGY4250. Distribution of EGFR amplification **(a)** and cell cycle status **(b)** amongst branches. Distribution of ChromVAR motif scores in branches for proneural motifs ASCL1 and OLIG2 **(c**,**d)**, ETV1 **(e)**, NFIX **(f)**, and mesenchymal motifs JUN::JUNB **(g)** and STAT3 **(h)**.

**Fig S12.**
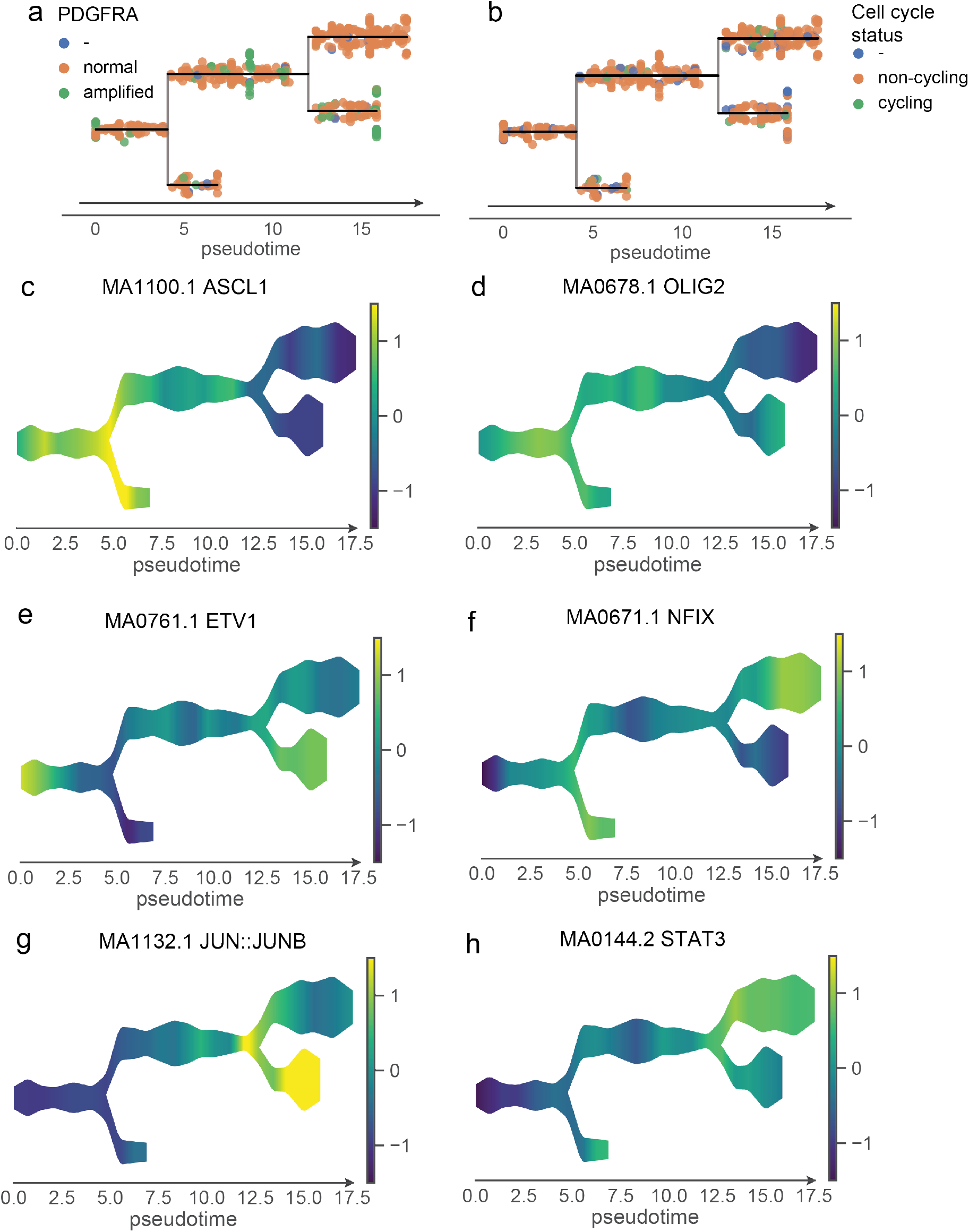
Pseudotime trajectory analysis of aGBM sample CGY4349. Distribution of PDGFRA amplification **(a)** and cycling status **(b)** amongst branches. Distribution of ChromVAR motif scores in branches for proneural motifs ASCL1 and OLIG2 **(c**,**d)**, ETV1 **(e)**, NFIX **(f)**, and mesenchymal motifs JUN::JUNB **(g)** and STAT3 **(h)**.

**Fig S13.**
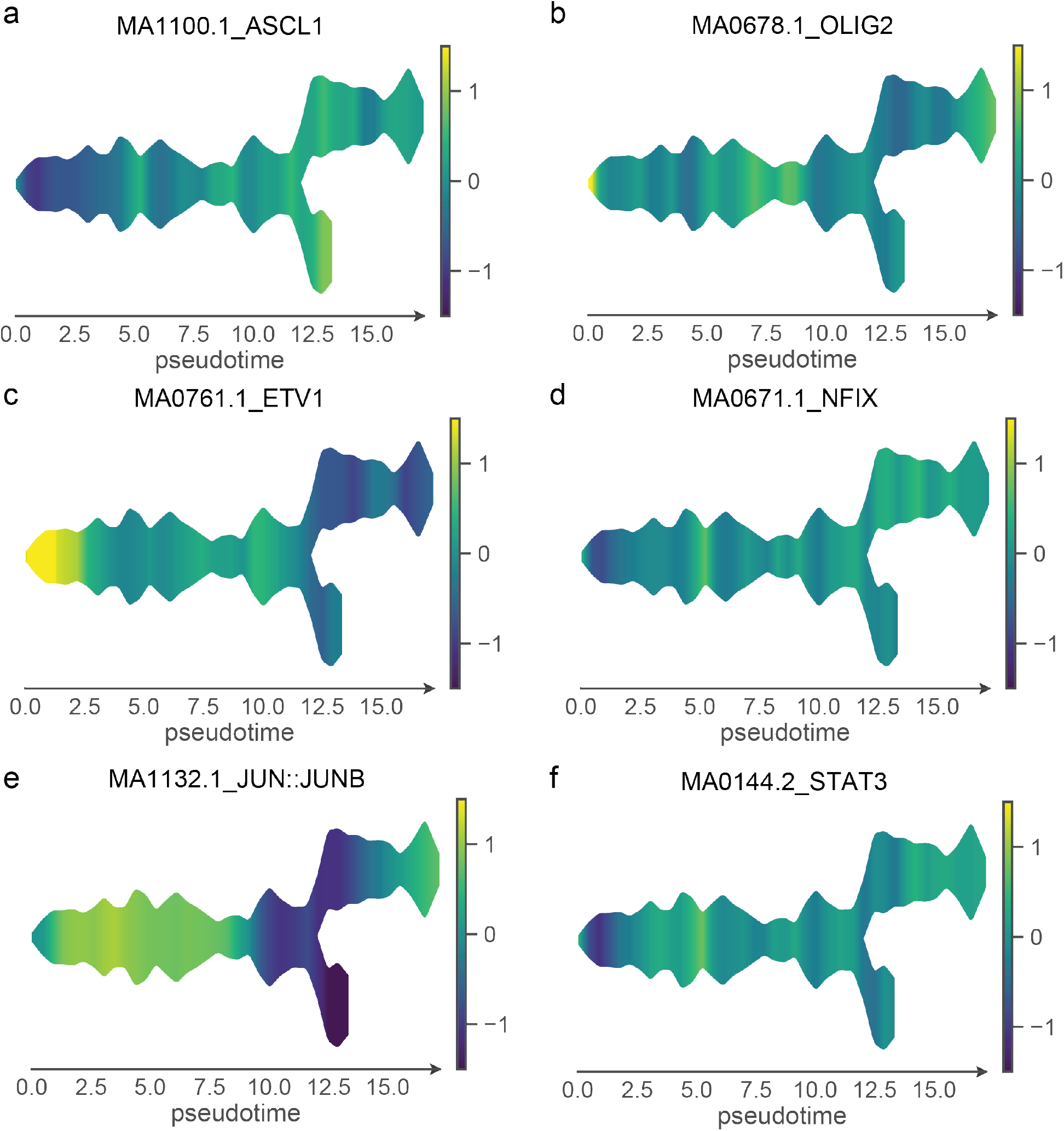
Pseudotime trajectory analysis of aGBM sample CGY4275. Distribution of ChromVAR motif scores in branches for proneural motifs ASCL1 and OLIG2 **(a**,**b)**, ETV1 **(c)**, NFIX **(d)**, and mesenchymal motifs JUN::JUNB **(e)** and STAT3 **(f)**.

## Notes

https://github.com/spcdot/CopyscAT

### Competing Interest Statement

The authors have declared no competing interest.

## References

1. B. Lim, Y. Lin, N. Navin, Advancing Cancer Research and Medicine with Single-Cell Genomics. Cancer Cell. 37, 456–470 (2020).

2. A. P. Patel, I. Tirosh, J. J. Trombetta, A. K. Shalek, S. M. Gillespie, H. Wakimoto, D. P. Cahill, B. V. Nahed, W. T. Curry, R. L. Martuza, D. N. Louis, O. Rozenblatt-Rosen, M. L. Suvà, A. Regev, B. E. Bernstein, Single-cell RNA-seq highlights intratumoral heterogeneity in primary glioblastoma. Science (80-.). (2014), doi:10.1126/science.1254257.

3. S. Darmanis, S. A. Sloan, D. Croote, M. Mignardi, S. Chernikova, P. Samghababi, Y. Zhang, N. Neff, M. Kowarsky, C. Caneda, G. Li, S. D. Chang, I. D. Connolly, Y. Li, B. A. Barres, M. H. Gephart, S. R. Quake, Single-Cell RNA-Seq Analysis of Infiltrating Neoplastic Cells at the Migrating Front of Human Glioblastoma. Cell Rep., 1399–1410 (2017).

4. J. Gojo, B. Englinger, L. Jiang, J. M. Hübner, M. L. Shaw, O. A. Hack, S. Madlener, D. Kirchhofer, Liu J. Pyrdol, V. Hovestadt, E. Mazzola, N. D. Mathewson, M. Trissal, D. Lötsch, C. Dorfer, C. Haberler, A. Halfmann, L. Mayr, A. Peyrl, R. Geyeregger, B. Schwalm, M. Mauermann, K. W. Pajtler, T. Milde, M. E. Shore, J. E. Geduldig, K. Pelton, T. Czech, O. Ashenberg, K. W. Wucherpfennig, O. Rozenblatt-Rosen, S. Alexandrescu, K. L. Ligon, S. M. Pfister, A. Regev, I. Slavc, W. Berger, M. L. Suvà, M. Kool, M. G. Filbin, Single-Cell RNA-Seq Reveals Cellular Hierarchies and Impaired Developmental Trajectories in Pediatric Ependymoma. Cancer Cell. 38, 44–59 (2020).

5. C. Neftel, J. Laffy, M. G. Filbin, T. Hara, M. E. Shore, G. J. Rahme, A. R. Richman, D. Silverbush, M. L. Shaw, C. M. Hebert, J. Dewitt, S. Gritsch, E. M. Perez, L. N. Gonzalez Castro, X. Lan, N. Druck, C. Rodman, D. Dionne, A. Kaplan, M. S. Bertalan, J. Small, K. Pelton, S. Becker, D. Bonal, Q.-D. Nguyen, R. L. Servis, J. M. Fung, R. Mylvaganam, L. Mayr, J. Gojo, C. Haberler, R. Geyeregger, T. Czech, I. Slavc, B. V. Nahed, W. T. Curry, B. S. Carter, H. Wakimoto, P. K. Brastianos, T. T. Batchelor, A. Stemmer-Rachamimov, M. Martinez-Lage, M. P. Frosch, I. Stamenkovic, N. Riggi, E. Rheinbay, M. Monje, O. Rozenblatt-Rosen, D. P. Cahill, A. P. Patel, T. Hunter, I. M. Verma, K. L. Ligon, D. N. Louis, A. Regev, B. E. Bernstein, I. Tirosh, M. L. Suvà, An Integrative Model of Cellular States, Plasticity, and Genetics for Glioblastoma. Cell. 178, 835–849.e21 (2019).

6. M. C. Vladoiu, I. El-Hamamy, L. K. Donovan, H. Farooq, B. L. Holgado, Y. Sundaravadanam, V. Ramaswamy, L. D. Hendrikse, S. Kumar, S. C. Mack, J. J. Y. Lee, V. Fong, K. Juraschka, D. Przelicki, A. Michealraj, P. Skowron, B. Luu, H. Suzuki, A. S. Morrissy, F. M. G. Cavalli, L. Garzia, C. Daniels, X. Wu, M. A. Qazi, S. K. Singh, J. A. Chan, M. A. Marra, D. Malkin, P. Dirks, L. Heisler, T. Pugh, K. Ng, F. Notta, E. M. Thompson, C. L. Kleinman, A. L. Joyner, N. Jabado, L. Stein, M. D. Taylor, Childhood cerebellar tumours mirror conserved fetal transcriptional programs. Nature. 572, 67–73 (2019).

7. I. Tirosh, A. S. Venteicher, C. Hebert, L. E. Escalante, A. P. Patel, K. Yizhak, J. M. Fisher, C. Rodman, C. Mount, M. G. Filbin, C. Neftel, N. Desai, J. Nyman, B. Izar, C. C. Luo, J. M. Francis, A. A. Patel, M. L. Onozato, N. Riggi, K. J. Livak, D. Gennert, R. Satija, B. V. Nahed, W. T. Curry, R. L. Martuza, R. Mylvaganam, A. J. Iafrate, M. P. Frosch, T. R. Golub, M. N. Rivera, G. Getz, O. Rozenblatt-Rosen, D. P. Cahill, M. Monje, B. E. Bernstein, D. N. Louis, A. Regev, M. L. Suvà, Single-cell RNA-seq supports a developmental hierarchy in human oligodendroglioma. Nature. 539, 309–313 (2016).

8. A. S. Venteicher, I. Tirosh, C. Hebert, K. Yizhak, C. Neftel, M. G. Filbin, V. Hovestadt, L. E. Escalante, M. L. Shaw, C. Rodman, S. M. Gillespie, D. Dionne, C. C. Luo, H. Ravichandran, R. Mylvaganam, C. Mount, M. L. Onozato, B. V. Nahed, H. Wakimoto, W. T. Curry, A. J. Iafrate, M. N. Rivera, M. P. Frosch, T. R. Golub, P. K. Brastianos, G. Getz, A. P. Patel, M. Monje, D. P. Cahill, O. Rozenblatt-Rosen, D. N. Louis, B. E. Bernstein, A. Regev, M. L. Suvà, Decoupling genetics, lineages, and microenvironment in IDH-mutant gliomas by single-cell RNA-seq. Science (80-.). 355 (2017), doi:10.1126/science.aai8478.

9. S. Müller, A. Cho, S. J. Liu, D. A. Lim, A. Diaz, CONICS integrates scRNA-seq with DNA sequencing to map gene expression to tumor sub-clones. Bioinformatics (2018), doi:10.1093/bioinformatics/bty316.

10. J. D. Buenrostro, P. G. Giresi, L. C. Zaba, H. Y. Chang, W. J. Greenleaf, Transposition of native chromatin for fast and sensitive epigenomic profiling of open chromatin, DNA-binding proteins and nucleosome position. Nat. Methods (2013), doi:10.1038/nmeth.2688.

11. J. D. Buenrostro, B. Wu, U. M. Litzenburger, D. Ruff, M. L. Gonzales, M. P. Snyder, H. Y. Chang, W. J. Greenleaf, Single-cell chromatin accessibility reveals principles of regulatory variation. Nature (2015), doi:10.1038/nature14590.

12. R. Killick, I. A. Eckley, Changepoint: An R package for changepoint analysis. J. Stat. Softw. (2014), doi:10.18637/jss.v058.i03.

13. M. Snuderl, L. Fazlollahi, L. P. Le, M. Nitta, B. H. Zhelyazkova, C. J. Davidson, S. Akhavanfard, D. P. Cahill, K. D. Aldape, R. A. Betensky, D. N. Louis, A. J. Iafrate, Mosaic amplification of multiple receptor tyrosine kinase genes in glioblastoma. Cancer Cell (2011), doi:10.1016/j.ccr.2011.11.005.

14. H. Chen, L. Albergante, J. Y. Hsu, C. A. Lareau, G. Lo Bosco, J. Guan, S. Zhou, A. N. Gorban, D. E. Bauer, M. J. Aryee, D. M. Langenau, A. Zinovyev, J. D. Buenrostro, G. C. Yuan, L. Pinello, Single-cell trajectories reconstruction, exploration and mapping of omics data with STREAM. Nat. Commun. 10, 1903 (2019).

15. A. N. Schep, B. Wu, J. D. Buenrostro, W. J. Greenleaf, ChromVAR: Inferring transcription-factor-associated accessibility from single-cell epigenomic data. Nat. Methods. 14, pages975–978 (2017).

16. A. P. Patel, I. Tirosh, J. J. Trombetta, A. K. Shalek, S. M. Gillespie, H. Wakimoto, D. P. Cahill, B. V. Nahed, W. T. Curry, R. L. Martuza, D. N. Louis, O. Rozenblatt-Rosen, M. L. Suvà, A. Regev, B. E. Bernstein, Single-cell RNA-seq highlights intratumoral heterogeneity in primary glioblastoma. Science. 344, 1396–1401 (2014).

17. M. D. Robinson, D. J. McCarthy, G. K. Smyth, edgeR: A Bioconductor package for differential expression analysis of digital gene expression data. Bioinformatics. 26, 139–140 (2010).

18. L. Scrucca, M. Fop, T. B. Murphy, A. E. Raftery, Mclust 5: Clustering, classification and density estimation using Gaussian finite mixture models. R J. 8, 289–317 (2016).

19. R. Killick, I. A. Eckley, Changepoint: An R package for changepoint analysis. J. Stat. Softw. 58 (2014), doi:10.18637/jss.v058.i03.

20. H. Chen, L. Albergante, J. Y. Hsu, C. A. Lareau, G. Lo Bosco, J. Guan, S. Zhou, A. N. Gorban, D. E. Bauer, M. J. Aryee, D. M. Langenau, A. Zinovyev, J. D. Buenrostro, G. C. Yuan, L. Pinello, Single-cell trajectories reconstruction, exploration and mapping of omics data with STREAM. Nat. Commun. (2019), doi:10.1038/s41467-019-09670-4.

21. A. N. Schep, B. Wu, J. D. Buenrostro, W. J. Greenleaf, ChromVAR: Inferring transcription-factor-associated accessibility from single-cell epigenomic data. Nat. Methods (2017), doi:10.1038/nmeth.4401.

22. A. Khan, O. Fornes, A. Stigliani, M. Gheorghe, J. A. Castro-Mondragon, R. Van Der Lee, A. Bessy, J. Chèneby, S. R. Kulkarni, G. Tan, D. Baranasic, D. J. Arenillas, A. Sandelin, K. Vandepoele, B. Lenhard, B. Ballester, W. W. Wasserman, F. Parcy, A. Mathelier, JASPAR 2018: Update of the open–access database of transcription factor binding profiles and its web framework. Nucleic Acids Res. (2018), doi:10.1093/nar/gkx1126.

23. H. Li, R. Durbin, Fast and accurate short read alignment with Burrows-Wheeler transform. Bioinformatics. 25, 1754–60 (2009).

24. H. Li, B. Handsaker, A. Wysoker, T. Fennell, J. Ruan, N. Homer, G. Marth, G. Abecasis, R. Durbin, The Sequence Alignment/Map format and SAMtools. Bioinformatics. 25, 2078–2079 (2009).

25. E. Talevich, A. H. Shain, T. Botton, B. C. Bastian, CNVkit: Genome-Wide Copy Number Detection and Visualization from Targeted DNA Sequencing. PLoS Comput. Biol. 12 (2016), doi:10.1371/journal.pcbi.1004873.

26. A. R. Quinlan, I. M. Hall, BEDTools: A flexible suite of utilities for comparing genomic features. Bioinformatics. 26, 841–2 (2010).

27. T. Stuart, A. Srivastava, C. Lareau, R. Satija, bioRxiv, in press, doi:10.1101/2020.11.09.373613.

28. A. Butler, P. Hoffman, P. Smibert, E. Papalexi, R. Satija, Integrating single-cell transcriptomic data across different conditions, technologies, and species. Nat. Biotechnol. 36, 411–420 (2018).

